# Spatially Resolved Transcriptomics Deconvolutes Histological Prognostic Subgroups in Patients with Colorectal Cancer and Synchronous Liver Metastases

**DOI:** 10.1101/2022.09.21.508569

**Authors:** Colin S Wood, Kathryn AF Pennel, Holly Leslie, Assya Legrini, Andrew J Cameron, Lydia Melissourgou-Syka, Jean A Quinn, Hester C van Wyk, Jennifer Hay, Antonia K Roseweir, Colin Nixon, Campbell SD Roxburgh, Donald C McMillan, Andrew V Biankin, Owen J Sansom, Paul G Horgan, Joanne Edwards, Colin W Steele, Nigel B Jamieson

**Author notes:** Corresponding Author: Nigel B Jamieson. Joint first/last authors (these authors contributed equally to this work) (Colin S Wood, Kathryn AF Pennel, Colin W Steele, Nigel B Jamieson).

## Abstract

**Background:** Patients demonstrating strong immune responses to primary colorectal cancer (CRC) have a survival benefit following surgery, while those with predominantly stromal microenvironments do poorly. Biomarkers to identify patients with colorectal cancer liver metastases (CRLM) who have good prognosis following surgery for oligometastatic disease remain elusive. The aim of this study was to determine the practical application of a simple histological assessment of immune cell infiltration and stromal content in predicting outcome following synchronous resection of primary CRC and CRLM, and to interrogate the underlying functional biology that drives disease progression.

**Methods:** Patients undergoing synchronous resection of primary CRC and CRLM underwent detailed histological assessment, panel genomic and bulk transcriptomic assessment, immunohistochemistry (IHC) and GeoMx Spatial Transcriptomics (ST) analysis. Integration with genomic features, pathway enrichment analysis and immune deconvolution were performed.

**Results:** High-immune metastases were associated with improved cancer specific survival (HR, 0.36, *P*=0.01). Bulk transcriptomic analysis was confounded by stromal content but ST demonstrated that the invasive edge of the metastases of long-term survivors was characterized by adaptive immune cell populations enriched for Type II Interferon signalling (NES=-2.05 *P.Adj*<0.005) and MHC-Class II Antigen Presentation (NES=-2.09 *P.Adj*<0.005). In contrast, patients with poor prognosis demonstrated increased abundance of regulatory T-cells and neutrophils with enrichment of Notch (NES=2.2 *P.Adj*=0.022) and TGF-β (NES=2.2 *P.Adj*=0.02) signalling pathways at the metastatic tumor centre.

**Conclusions:** Histological assessment stratifies outcome in patients undergoing synchronous resection of CRLM. ST analysis reveals significant intra-tumoral and inter-lesional heterogeneity with underlying transcriptomic programmes identified in driving each phenotype.

**TRANSLATIONAL RELEVANCE:** The current study demonstrates that accurate histological assessment of immune cell infiltration and stromal content can define survival in patients following resection of oligometastatic liver disease when presenting synchronously with primary colorectal cancer. A spatial transcriptomic approach has demonstrated heterogeneity between patients, between matched lesions in the same patient and within individual lesions. Patients with high immune infiltrates at the invasive margin demonstrated lymphocytic infiltration and associated upregulated adaptive immune pathways in long term survivors. In specimens with low immune infiltrate at the tumor edge a significant reduction in survival was observed, this was determined by upregulated immunosuppressive pathways and a predominance of innate immune cells surrounding metastases. Spatial transcriptomics can be used to examine drivers of metastatic progression in CRC and identifies patients with reactive and suppressed immune microenvironments. Application across a larger cohort will build the cartography of CRLM, while in future, studies may assess application of this technology to pre and post treatment biopsy samples with the aim of predicting individual therapeutic responses. The current study has highlighted discrepancies between bulk and ST derived data whilst demonstrating accuracy of deconvoluted transcriptome to determine immune profiling. Now that ST strategies are becoming more achievable at scale, this has implications for the interpretation of the bulk transcriptomic signatures both of primary and metastatic CRC.

## INTRODUCTION

Colorectal Liver Metastases (CRLM) remain the largest contributor to death for patients with Colorectal Cancer (CRC), with approximately 50% experiencing disease recurrence at this site (1). Modern surgical techniques including portal vein embolization and parenchyma sparing surgery have increased the number of patients eligible for resection, however recurrence post-metastastectomy approaches 70% and intuitively, patients who recur have poorer outcomes (2, 3). To better predict outcome following CRLM resection, Fong et al. derived a score based on clinicopathological factors including nodal burden, synchronicity of presentation, number of CRLM and CEA level (4). More recently genomic profiling has identified *KRAS*, *BRAF* and *KRAS*/*TP53* co-mutations associating with poor prognosis (5–7), however, patient responses to treatment are unpredictable and successful development of novel therapies targeting stage IV CRC remain challenging (8). Expanding therapeutic targeting beyond mutational status to immune landscape and transcriptomic phenotypes may hold insight into the future management of CRLM (9).

Assessment of the tumor microenvironment (TME) in primary CRC, most notably the Immunoscore to determine the CD8/CD3+ T cell ratio, has become a validated prognostic tool (10, 11). Simpler, yet powerful metrics evaluating Haematoxylin and Eosin (H&E) stained sections include Klintrup-Mäkinen (KM) grade, assessing immune cell density at tumor invasive edge (IE) and Tumor Stroma Percentage (TSP) measuring stromal density. Both have been applied to primary CRC, however, their prognostic utility in the metastatic setting remains unexplored, with biological interrogation also lacking (12, 13).

Four Consensus Molecular Subtypes (CMS) of primary CRC have been defined through bulk transcriptomic profiling generating a roadmap for translational CRC biology (14) but have failed to project into the metastatic setting (15). Increasingly, constraints of bulk transcriptomics are apparent with ‘stromal noise’ potentially concealing biological pathway discovery and intra-tumoral heterogeneity characterisation (16, 17). While single-cell RNA sequencing has led to important discoveries (18, 19), crucially, topographic cellular orientation is lost along with vital biological insights relating to tissue morphology, cellular interactions and TME location-specific cellular expression. To overcome these limitations, spatial transcriptomics (ST) solutions aim to provide rich molecular insight while maintaining histological architecture. The Nanostring© GeoMx Digital Spatial Profiler (DSP) is a novel ST platform enabling hi-plex, high-throughput characterisation of user defined regions on Formalin-Fixed Paraffin-Embedded (FFPE) tissue (20) employing UV-photocleavable barcodes hybridized to multiplex immuno-fluorescence (mIF) stained tissue enabling up to whole transcriptome analysis.

To better understand the mechanisms underlying metastatic development, the current study sought to investigate the prognostic utility of basic immunological assessments (KM and TSP) in a cohort of synchronously resected primary CRC with matched CRLM, integrating detailed immunohistochemistry (IHC), genome panel and bulk transcriptomic data to clearly define patient groups. Furthermore, an exploratory spatial transcriptomic assessment using the Nanostring GeoMx DSP platform to interrogate the functional biology underlying clinically relevant subtypes in matched primary and CRLM to identify the features of disease progression was performed.

## MATERIALS AND METHODS

### Cohort Characteristics

41 patients undergoing synchronous resection of primary CRC and CRLM with curative intent between April 2002 and June 2010 at Glasgow Royal Infirmary by a single surgeon (PH) were included (Table 1). Patients were identified from a prospectively maintained database and represent a consecutive cohort of resected patients with mature 10-year postoperative follow-up including recurrence and mortality data. Patients were excluded if pathology slides were unavailable or if survival data was incomplete. Application to access patient tissue was authorized by the NHS Greater Glasgow and Clyde Biorepository under their NHS Research Ethics Committee (REC) approval with ethical approval granted in biorepository application #357, West of Scotland Ethics 22/WS/0207. Patients were followed up in the postoperative period at 1 month, then 6-monthly until 2 years, and thereafter annually until 5 years, at which time-point they were discharged from follow-up. Date and cause of death was confirmed via access to the NHS Greater Glasgow and Clyde Clinical Portal. Records were complete until 19^th^ November 2020, which served as the censor date. Cancer-specific survival (CSS) was measured from the date of surgery until the date of death from CRC.

**Table 1:**
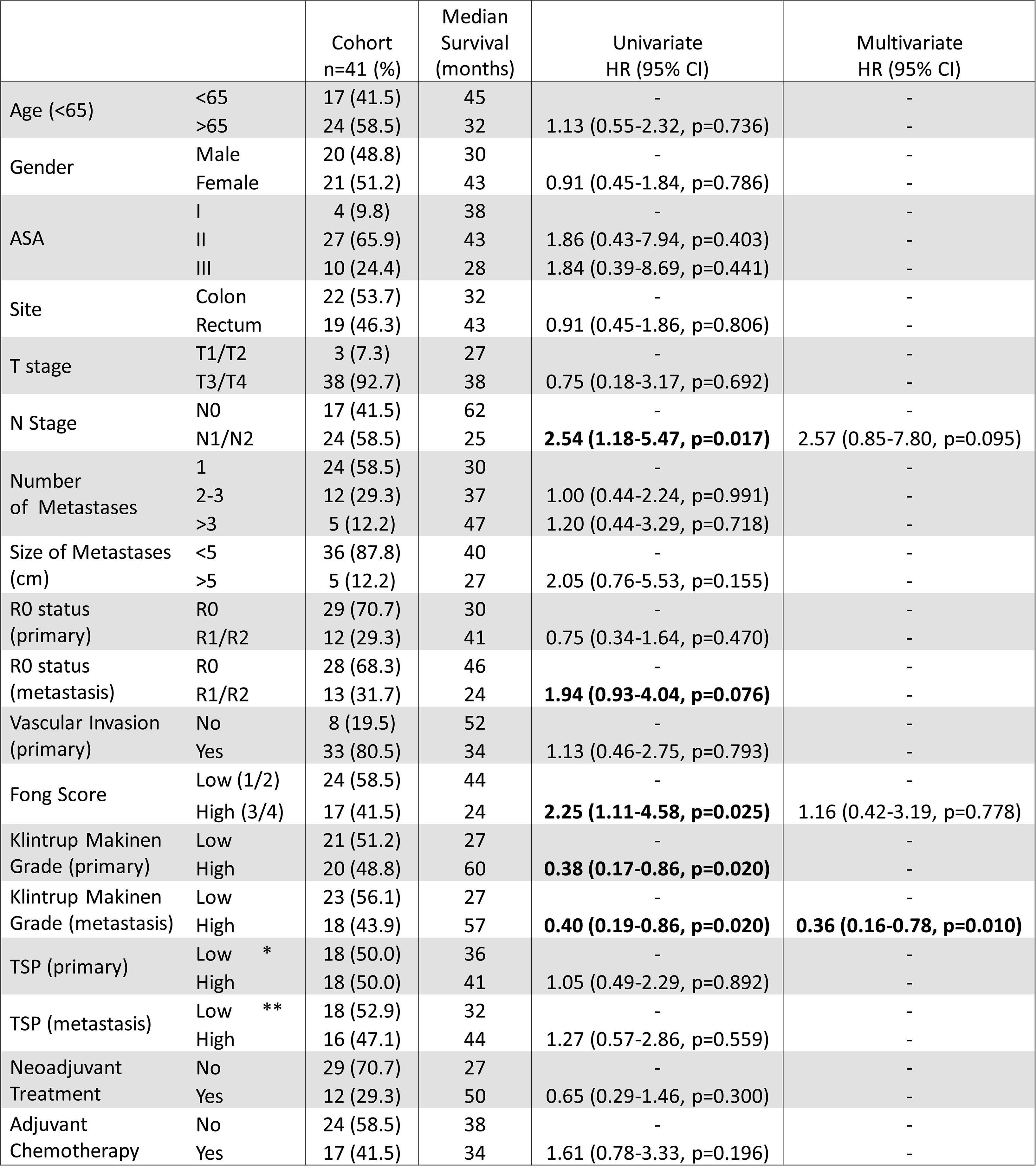
Clinicopathological, morphological and treatment characteristics for synchronously resected primary CRC and paired CRLM. Clinicopathological and treatment data for 41 patients managed by synchronous resection of primary CRC and CRLM are displayed. Univariate and multivariate survival analysis presented and calculated using adjusted Cox proportional hazards regression model. * 36 primary CRC assessed for TSP ** 34 CRLM assessed for TSP CI, confidence interval; HR, hazard ratio; ASA, American Society of Anesthesiologists Physical Status Classification; TSP: Tumour Stromal Percentage

|  |  | Cohort<br>n=41 (%) | Median<br>Survival<br>(months) | Univariate<br>HR (95% CI) | Multivariate<br>HR (95% CI) |
| --- | --- | --- | --- | --- | --- |
| Age (<65) | <65 | 17 (41.5) | 45 | - | - |
|  | >65 | 24 (58.5) | 32 | 1.13 (0.55-2.32, p=0.736) | - |
| Gender | Male | 20 (48.8) | 30 | - | - |
|  | Female | 21 (51.2) | 43 | 0.91 (0.45-1.84, p=0.786) | - |
| ASA | I | 4 (9.8) | 38 | - | - |
|  | II | 27 (65.9) | 43 | 1.86 (0.43-7.94, p=0.403) | - |
|  | III | 10 (24.4) | 28 | 1.84 (0.39-8.69, p=0.441) | - |
| Site | Colon | 22 (53.7) | 32 | - | - |
|  | Rectum | 19 (46.3) | 43 | 0.91 (0.45-1.86, p=0.806) | - |
| T stage | T1/T2 | 3 (7.3) | 27 | - | - |
|  | T3/T4 | 38 (92.7) | 38 | 0.75 (0.18-3.17, p=0.692) | - |
| N Stage | N0 | 17 (41.5) | 62 | - | - |
|  | N1/N2 | 24 (58.5) | 25 | **2.54 (1.18-5.47, p=0.017)** | 2.57 (0.85-7.80, p=0.095) |
| Number<br>of Metastases | 1 | 24 (58.5) | 30 | - | - |
|  | 2-3 | 12 (29.3) | 37 | 1.00 (0.44-2.24, p=0.991) | - |
|  | >3 | 5 (12.2) | 47 | 1.20 (0.44-3.29, p=0.718) | - |
| Size of Metastases<br>(cm) | <5 | 36 (87.8) | 40 | - | - |
|  | >5 | 5 (12.2) | 27 | 2.05 (0.76-5.53, p=0.155) | - |
| R0 status<br>(primary) | R0 | 29 (70.7) | 30 | - | - |
|  | R1/R2 | 12 (29.3) | 41 | 0.75 (0.34-1.64, p=0.470) | - |
| R0 status<br>(metastasis) | R0 | 28 (68.3) | 46 | - | - |
|  | R1/R2 | 13 (31.7) | 24 | **1.94 (0.93-4.04, p=0.076)** | - |
| Vascular Invasion<br>(primary) | No | 8 (19.5) | 52 | - | - |
|  | Yes | 33 (80.5) | 34 | 1.13 (0.46-2.75, p=0.793) | - |
| Fong Score | Low (1/2) | 24 (58.5) | 44 | - | - |
|  | High (3/4) | 17 (41.5) | 24 | **2.25 (1.11-4.58, p=0.025)** | 1.16 (0.42-3.19, p=0.778) |
| Klintrup Makinen<br>Grade (primary) | Low | 21 (51.2) | 27 | - | - |
|  | High | 20 (48.8) | 60 | **0.38 (0.17-0.86, p=0.020)** | - |
| Klintrup Makinen<br>Grade (metastasis) | Low | 23 (56.1) | 27 | - | - |
|  | High | 18 (43.9) | 57 | **0.40 (0.19-0.86, p=0.020)** | **0.36 (0.16-0.78, p=0.010)** |
| TSP (primary) | Low * | 18 (50.0) | 36 | - | - |
|  | High | 18 (50.0) | 41 | 1.05 (0.49-2.29, p=0.892) | - |
| TSP (metastasis) | Low ** | 18 (52.9) | 32 | - | - |
|  | High | 16 (47.1) | 44 | 1.27 (0.57-2.86, p=0.559) | - |
| Neoadjuvant<br>Treatment | No | 29 (70.7) | 27 | - | - |
|  | Yes | 12 (29.3) | 50 | 0.65 (0.29-1.46, p=0.300) | - |
| Adjuvant<br>Chemotherapy | No | 24 (58.5) | 38 | - | - |
|  | Yes | 17 (41.5) | 34 | 1.61 (0.78-3.33, p=0.196) | - |

### Immune landscape morphology analysis

Synchronously resected CRC and CRLM FFPE tissues were sectioned, stained with H&E, and scored using previously described methods (Supplemental Figure 1) (12, 21). KM grading of immune cell infiltration was scored from 0-3 for the depth of immune cells at the tumor invasive edge (IE) according to appearances at the deepest area of tumor invasion. Score 0 - no increase in inflammatory cells at the deepest point of invasive margin; 1 - mild and patchy increase in inflammatory cells; 2 - a prominent inflammatory reaction forming a band at the invasive margin with some evidence of destruction of cancer cell islands; 3 - a florid cup-like inflammatory infiltrate at the IE with frequent destruction of cancer cells. Tumors scored 2-3 were assigned KM^high^ with the remainder classified as KM^low^. For TSP evaluation, tumors with > 50% stroma were classified as stroma^high^, with the remaining allocated as stroma^low^ (13). 15 primary CRC and CRLM were co-scored and in keeping with previously reported low inter-observer variability (21), the average correlation of both scoring systems was 0.87 (high) between observers.

### GPOL panel Mutational analysis

DNA was extracted from FFPE sections and standardized to a concentration of 4ng/µl using the DNeasy kit (Qiagen). Mutational landscaping was performed by the Glasgow Precision Oncology Laboratory employing an in-house Genomic panel assay of 151 cancer-associated genes (Supplemental Table 1) (22). Targeted capture libraries were prepared from 150-200ng DNA. Sequencing was performed using an Illumina® HiSeq400 (San Diego, CA, USA). The *maftools* (23) package was used to generate oncoplots, forest plots and co-barplots to compare the mutational landscape for primary CRC and CRLM with morphological and immune cell integration.

### Immunohistochemistry

Detailed staining methods for CD3 and CD66b are included in Supplemental Methods. Visualisation employed the SlidePath digital image hub (Leica Biosystems, Milton Keynes, UK) using a Hamamatsu NanoZoomer (Welwyn Garden City, Hertfordshire, UK). QuPath was used for image analysis (Version: 0.3.2, University of Edinburgh, UK) (24). From each primary and CRLM, 4 rectangular regions (mean perimeter 4740µm, mean area 1.38mm^2^) corresponding to the IE and tumor centre (TC) were annotated and positive cell detection was performed using in-built QuPath functionality. An R Studio pipeline was constructed (v1.2.1335 R Studio, Boston, USA) to compare the number of positive cell detections per mm^2^ with the median cell count used to determine high and low cut-off for categorical variables.

### Nanostring nCounter PanCancer IO360 Bulk Transcriptomic Panel

Macrodissection of tumor regions including TME and epithelium from unstained 10um thick sections was guided by an H&E image (see Supplemental Methods). RNA was extracted using the AllPrep DNA/RNA FFPE Kit (Qiagen) according to the manufacturer’s protocol, using xylene for deparaffinisation (average RNA concentration = 35ng/ul). RNA quantity was assessed using RNA BR assay and the Qubit^®^ 2.0 fluorometer (Invitrogen, Life Technologies). RNA integrity (RIN) values were determined using Agilent 2100 Bioanalyzer (Agilent Technologies) (maximum RIN = 2.1). Gene expression analysis was performed using the Nanostring nCounter IO360 panel (770 genes). Data acquisition was performed by using Nanostring’s Digital Analyser (FOV, 555). Raw gene expression count data were normalized using NanoString nSolver 4.0 software using 6 positive controls and 8 negative controls to account for background noise and sample variation across runs (GeNorm Algorithm).

### Nanostring GeoMx Digital Spatial Profiling

The DSP protocol has been described by Merritt et al (20) and consists of slide preparation including antigen retrieval and staining with immunofluorescent markers, hybridisation of tissue with UV-photocleavable probes, scanning, region selection, probe collection, library preparation and sequencing. Supplemental Methods provides detailed description of slide preparation. Four channels are available for the detection of four customisable morphology markers: FITC/525nm, Cy3/568nm, Texas Red/615nm, and Cy5/666nm (20). One channel is reserved for the nuclear stain (DAPI). In this experiment, Pan-Cytokeratin (PanCK), CD45 for global immune cell population and α-SMA for fibroblasts and collagenous architecture constituted the other channels. The primary CRC and paired CRLMs were mounted on the same slide (4 matched pairs on 4 slides) prior to hybridization with the Cancer Transcriptome Atlas (CTA) panel of probes corresponding to 1,825 genes (Nanostring). The CTA panel is designed to comprehensively characterize immune activity and tumor biology within the TME (27).

Following hybridisation, the slides were scanned on the GeoMx instrument with the workspace used to select regions of interest (ROI). In CRLM, distinct regions at the IE were observed, those which stained strongly for CD45 were annotated as “metastatic Invasive Edge - immune ” (mIE) (pink). ROIs at the IE of metastases that stained more prominently for αSMA were annotated as “metastatic Invasive Edge - stroma” (mSE) (yellow). PanCK stained epithelial regions in the centre of CRLM were annotated as “metastatic Tumor Centre” (mTC) (blue). In primary CRC, central epithelial regions were annotated as “primary Tumor Centre” (pTC) (dark green) and regions at the IE where there was a clear interface between epithelium and TME comprising clear immune cell populations were annotated as “primary Invasive Edge” (pIE) (light green). 3 of the primary lesions had Tertiary Lymphoid Regions (pTLR) (brown). One TLR was identified within metastasis (mTLR) (orange). Within each primary and CRLM, ROIs were selected with a range of mTC, mIE, mSE and mTLR (Supplemental Figures 2-5). Each of the 48 circular ROIs were 500µm diameter (Area 196795µm^2^) with average nuclei count of 1371 (range 707-2055).

**Figure 1.**
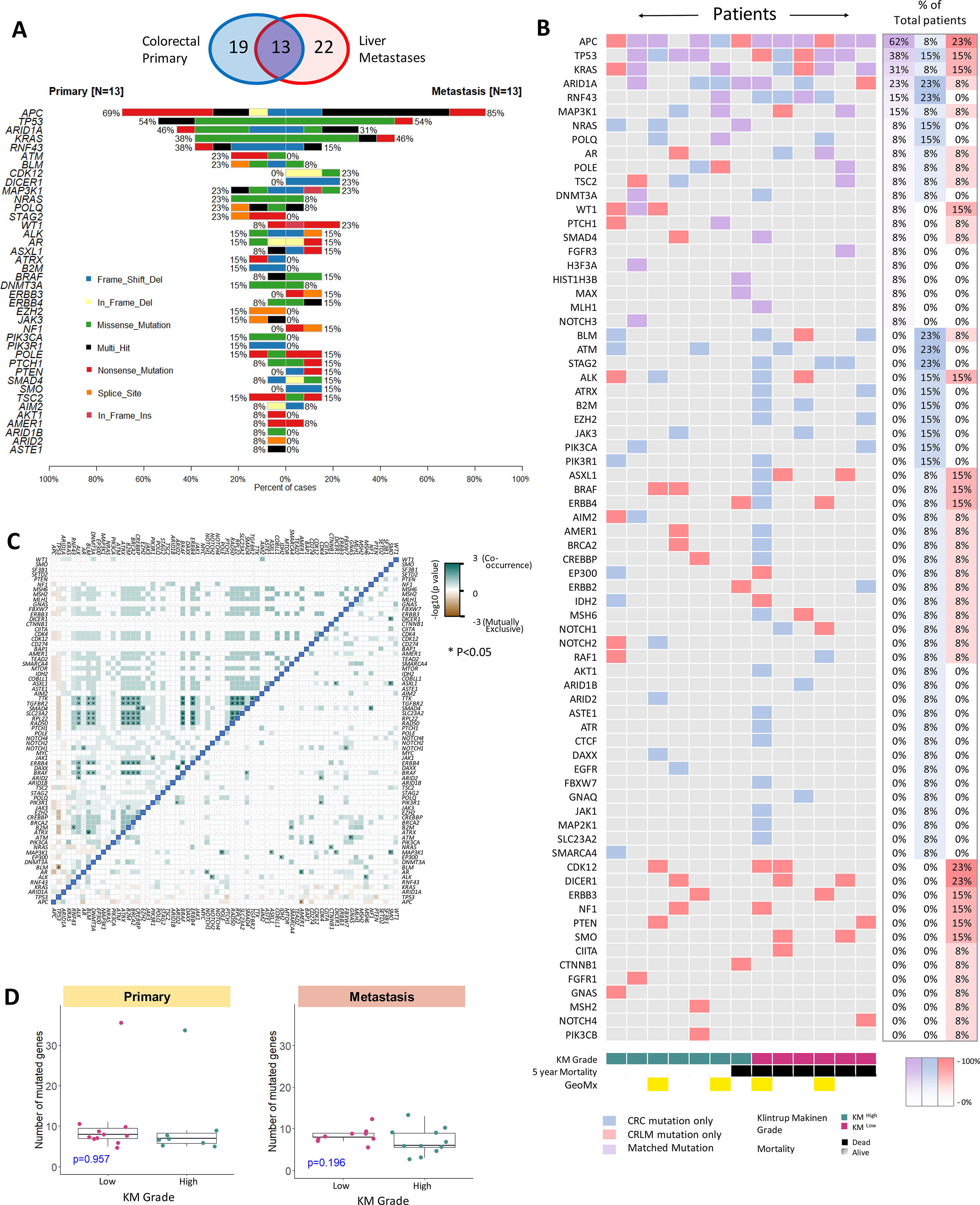
Mutational Characterisation of primary CRC and CRLM. A: Venn diagram demonstrating primary CRC and CRLM that underwent genomic analysis. Co-Barplot illustrating most frequently mutated genes across 13 matched primary CRC and CRLM including mutation type. Genes are ordered by mutational frequency. Sections were sequenced using GPOL (Glasgow Precision Oncology Laboratory) mutational panel. B: Oncoplot demonstrating concurrent mutations in the 13 matched lesions. Patients are ranked according to co-mutational burden on the y-axis and ranked according to KM grade on the x-axis. Blue denotes gene mutated in primary only, red denotes gene mutated in metastasis only, purple denotes mutated in primary and metastasis. The right hand 3 columns denote percentage of total patients with each mutation type. C: Correlation matrix demonstrating co-occurrence of mutations with left of the blue demarcation line representing primary CRC, right of the blue demarcation line representing CRLM (pair-wise Fisher’s Exact test *P < 0.05). Gene names are displayed along the x and y-axis ordered by mutational frequency. Dark green boxes represent significant co-occurrence D: Box plot illustrating mutational burden in primary CRC and CRLM according to KM grade using the Mann Whitney test to assess for statistically significant difference between KM groups

**Figure 2.**
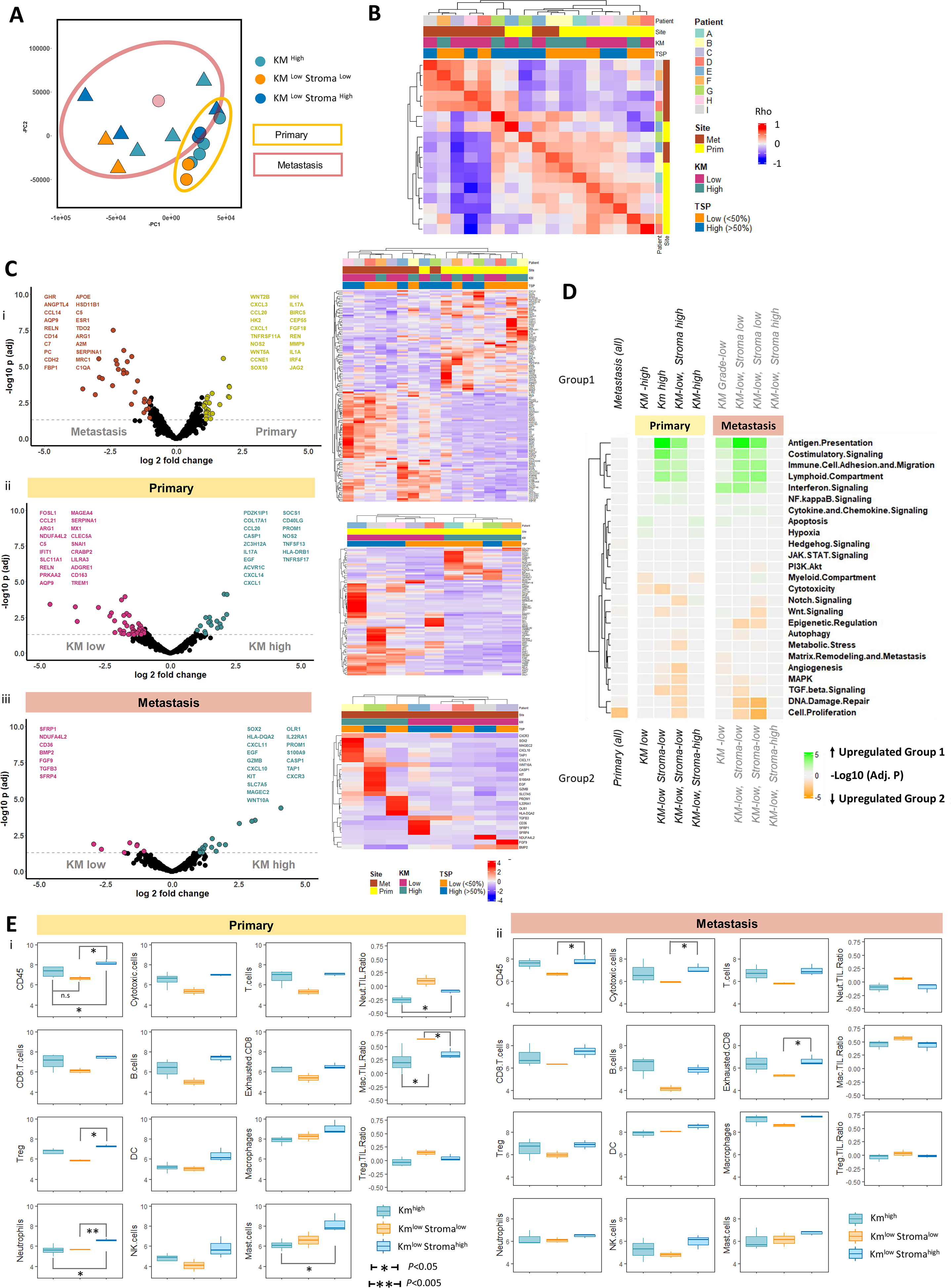
BULK IO360 transcriptomic characterisation of matched primary CRC and CRLM. A: PCA plot demonstrating 2 principal components of minimal variance for all samples. Primary CRC samples are demonstrated by circles and yellow outline. CRLM are demonstrated by triangular points and brown outline. KM^high^, KM^low^ Stroma^low^ and KM^low^ Stroma^high^ samples are depicted by colour. B: Unsupervised analysis using gene expression correlation matrix for all samples. Patient, site, KM grade, tumor stromal percentage (TSP) are depicted by key. Spearman correlation of all expressed genes performed between each sample sequenced and plotted on the heatmap. k-means clustering of heatmap to demonstrate correlated samples. Red represents strong correlation. Blue represents negative correlation. C: Volcano plots demonstrating differential gene expression results and clustered heatmap of significant genes for i) All primary CRC vs CRLM ii) KM grade: KM ^high^ vs KM ^low^ primary CRC iii) KM grade: KM ^high^ vs KM ^low^ CRLM. X-axis of volcano plot demonstrates log2fold change, y-axis demonstrates –log10*P*. Coloured points demonstrate significant changes in gene expression between groups (*P*<0.05 and logFC>1.5). Volcano plots demonstrate top 20 differentially expressed genes for each group. D: Heatmap demonstrating Gene Set Enrichment Analysis (GSEA) results comparing the different tumors grouped according to KM grade and TSP using io360 curated gene sets annotated on the right of the diagram. Heatmap squares represent Log10 Adjusted P-value. Green demonstrates upregulation in group 1, orange represents upregulation in group 2. The heatmap is clustered by y-axis only to demonstrate frequently upregulated gene sets. E: Boxplot comparison of immune cell populations and selected cell:cell ratios between KM grade and TSP segregated groups using deconvolution software included in the nCounter package. Annotated subgroups are: Km^high^, Km^low^ Stroma^low^, Km^low^ Stroma^high^. Y-axis represents log10 of estimated cell count. Primary CRC are represented in (i) and CRLM in (ii).

Once ROI selection was complete, collection was initiated whereby photo-cleavable oligonucleotide probes in the ROIs were exposed to UV-light to cleave the UV-sensitive probes. The released probe-specific DSP barcodes were then aspirated from selected ROIs and collected into the 96-well DSP plate. The probes, following rehydration with DEPC-treated water, were then added to the corresponding well of a new 96-well PCR plate containing the GeoMx Seq Code primers and the PCR Master Mix. Details of library preparation and sequencing are included in Supplemental Methods.

### Survival Analysis

All analyses using RStudio utilized v1.2.1335 of RStudio (R build version 4.1.1). Kaplan-Meier survival analysis curves (Log-Rank test) were generated using *survival* (25) and *survminer* (26) packages. Cox-regression analysis was performed using *finalfit* (27) and *hmisc* (28) packages. Statistical significance was set to *P* < 0.05 unless otherwise stated.

### IO360 Panel and GeoMx Spatial Transcriptomic analysis

IO360 panel analysis data underwent QC and normalisation in the proprietary nCounter Advanced Analysis Suite. For the GeoMx data, the Digital Count Conversion (DCC) files were uploaded onto the GeoMx DSP analysis suite (Nanostring), where they underwent quality control (QC), filtering, Q3 normalisation, and background correction. The normalized counts from each were downloaded into RStudio. Principal Component Analysis (PCA) was performed using *prcomp* (base R) and plotted using *ggplot2* (29). Differential Gene Expression (DGE) was performed using the Exact Test as part of edgeR package (30). Volcano plots were generated using *ggplot2* (29). Heat maps were generated using *ComplexHeatMap* (31). Gene set enrichment analysis (GSEA) was performed using the *fgsea* package (32). Single Sample Gene Set Enrichment Analysis (ssGSEA) was performed using the *GSVA* package (33). *ClusterProfiler* (34) was used to interrogate the Reactome curated database (35). Immune cell spatial deconvolution for nCounter IO360 data was performed in the nCounter Advanced Analysis suite with RStudio employed for analysis and visualisation. Immune spatial deconvolution of GeoMx derived CTA data was performed using the Bioconductor *SpatialDecon* tool (36).

## RESULTS

### Clinicopathological and morphological characteristics determine patient outcome

Baseline clinicopathological and treatment details for the 41 patients with synchronously resected CRC and CRLM are described in Table 1. 59% were ≥65 years old with similar gender distribution. Both rectal (46%) and colonic (54%) primary CRC were included, with the majority being stage T3 or T4 (93%), 59% had lymph node metastases and 29% received neoadjuvant chemotherapy. The 5 and 10-year mortality rate was 64% and 82% respectively.

The impact of traditional clinicopathological variables on outcome was assessed (Table 1). N-Stage (HR, 2.54, *P*=0.017) and Fong Score (HR, 2.25, *P*=0.025) were prognostic on univariate analysis. Gross morphological immune and stromal feature evaluation demonstrated KM^high^ in the primary CRC (HR, 0.38, *P*=0.02) and CRLM (HR, 0.40, *P*=0.02) predicted CSS (Table 1). In a multivariate model, KM^high^ within CRLM was the most significant prognostic factor (HR, 0.36, *P*=0.01).

### Genomic Characterisation of Synchronous CRLM

Mutational analysis was performed to determine if genomic landscape underpins immune morphology in CRC and paired CRLM. Gene mutation panel profiling for 19 primary CRC and 22 CRLM was performed, of which 13 were matched pairs (Figure 1A). For the most frequently mutated genes, a similar mutation rate was noted between primary CRC and CRLM (Figure 1A): *APC* 69% primary CRC and 85% CRLM; *TP53* 54% and 54%; *ARID1A* 46% and 31% and *KRAS* 38% and 46%. These genes were also the most frequently co-mutated genes between paired sites (Figure 1B). Out-with the frequently occurring mutations, rarer mutations were less likely to be concordant between sites. There were no significantly discordant genes between primary CRC and CRLM (Supplemental Figure 6A), however, *CDK12*, *ERBB3* and *DICER1* were exclusively mutated in CRLM (Supplemental Figure 6A). *APC*, *TP53*, *ARID1A* and *KRAS* mutations were mutually exclusive of other mutations in both primary CRC and CRLM (Figure 1C). Multiple clusters of co-occurring mutations were noted including *BRAF*, *ERBB4*, *BRCA2* and *TGFBR2* which associated with two patients with hyper-mutation status (>10 mutations). These were present in primary CRC only suggesting the hyper-mutated state was not conserved between primary and CRLM.

While we expected tumor mutational burden to associate with immune morphology, neither KM grade nor TSP were associated with mutational frequency or landscape (Figure 1D). *KRAS*, *TP53* co-mutation was more common in CRLM (6) than CRC (3). Of the patients with *KRAS, TP53* CRLM co-mutation, 5 died early following surgery (range 20 - 42 months) whilst only one of these patients survived long term (Log-rank, *P*=0.26, Supplemental Figure 6B). No other genomic subgroups associated with prognosis.

### Bulk transcriptomic analysis of prognostic subtypes

To gain insight into the gene expression profile associated with immune morphology subtypes, 9 matched primary CRC and CRLM were selected for nCounter bulk transcriptomic IO360 panel analysis. (Supplemental Figure 7). PCA plot (Figure 2A) demonstrates clear separation of primary CRC (N = 9) and CRLM (N = 8) with wide inter-lesional discrimination according to KM and TSP status evident for both primary and metastases (one CRLM failed QC).

An unsupervized hierarchical clustering correlation matrix confirmed tumor site was the predominant determinant of transcriptomic profile (Figure 2B). When segregated into primary CRC and CRLM groups, KM status further discriminates gene expression at both sites, particularly within CRLM (Supplemental Figure 8A and B). These gene lists were defined by DGE and predictably the highest number of differentially expressed genes resulted from primary CRC and CRLM comparison (Figure 2C). Between DGE of KM^high^ and KM^low^ in primary and CRLM, there was minimal overlap. An apparent stromal subgroup was apparent within KM^low^ lesions (Figure 2C).

GSEA comparing primary CRC and CRLM demonstrated a single significantly upregulated pathway in primary CRC (Cell Proliferation, normalized enrichment score (NES)=2.05, *P.Adj*=0.002) (Figure 2D). KM^low^ lesions were separated into KM^low^ Stroma^low^ and KM^low^ Stroma^high^ groups according to the PCA plot (Figure 2A). When pairwise comparison of each group was performed using GSEA and plotted in a clustered heatmap, concordant immune pathways enrichment for KM^high^ and KM^low^ Stroma^high^ tumors was demonstrated in both primary CRC and CRLM including Antigen Presentation, Costimulatory Signalling, Immune Cell Adhesion and Migration, Lymphoid Compartment and Interferon Signalling (Figure 2D). These data suggest that despite significant prognostic and histological differences, KM^high^ and KM^low^ Stroma^high^ lesions demonstrate similar bulk transcriptomic immune pathway dysregulation. In contrast, KM^low^ Stroma^low^ lesions were characterized by aberrations of Cell Proliferation, DNA Damage Repair and TGF-β Signalling at both primary and metastatic site.

Immune cell deconvolution of the IO360 panel transcriptome data demonstrated that while T-cell and B-cell abundance was similar for KM^high^ and KM^low^ Stroma^high^ lesions across primary CRC and CRLM, in KM^low^ Stroma^low^ lesions these populations were less prevalent (Figure 2E). Differences were less notable in myeloid derived cell populations (macrophages, neutrophils and NK cells). To highlight these differences in cell populations, immune cell ratios were calculated demonstrating that neutrophil: T-cell, macrophage: T-cell and NK: T-cell ratios were highest in KM^low^ Stroma^low^ lesions, most significantly in primary CRC (Figure 2E). These data may suggest high tumor stroma content disproportionately confounding the bulk transcriptomic signal.

### Immune cell characterisation defines KM phenotype

IHC was performed to further delineate the role of specific immune cells in prognostic subtypes. Samples were stained for CD3 (T-lymphocytes) (primary n=21; CRLM n=28) and CD66b (granulocytes/neutrophils) (primary n=21; CRLM n=29) based on tumorigenic role of neutrophils demonstrated in a pre-clinical model by our group (37). Representative images are shown in Figure 3A. Increased CD3 density was demonstrated at the CRLM IE compared with primary CRC IE (median 683.5, Interquartile Range (IQR): 272-1039 vs median 1142.2, IQR: 810-1702, *P*=0.009) (Figure 3B). Conversely, CD3 density was reduced in the CRLM TC compared with primary CRC TC (median 190, IQR: 71-273 vs median 443, IQR: 150-629, *P*=0.006) (Figure 3B). CD66b count in contrast did not differ significantly between primary and metastasis at TC or IE (Figure 3B).

**Figure 3:**
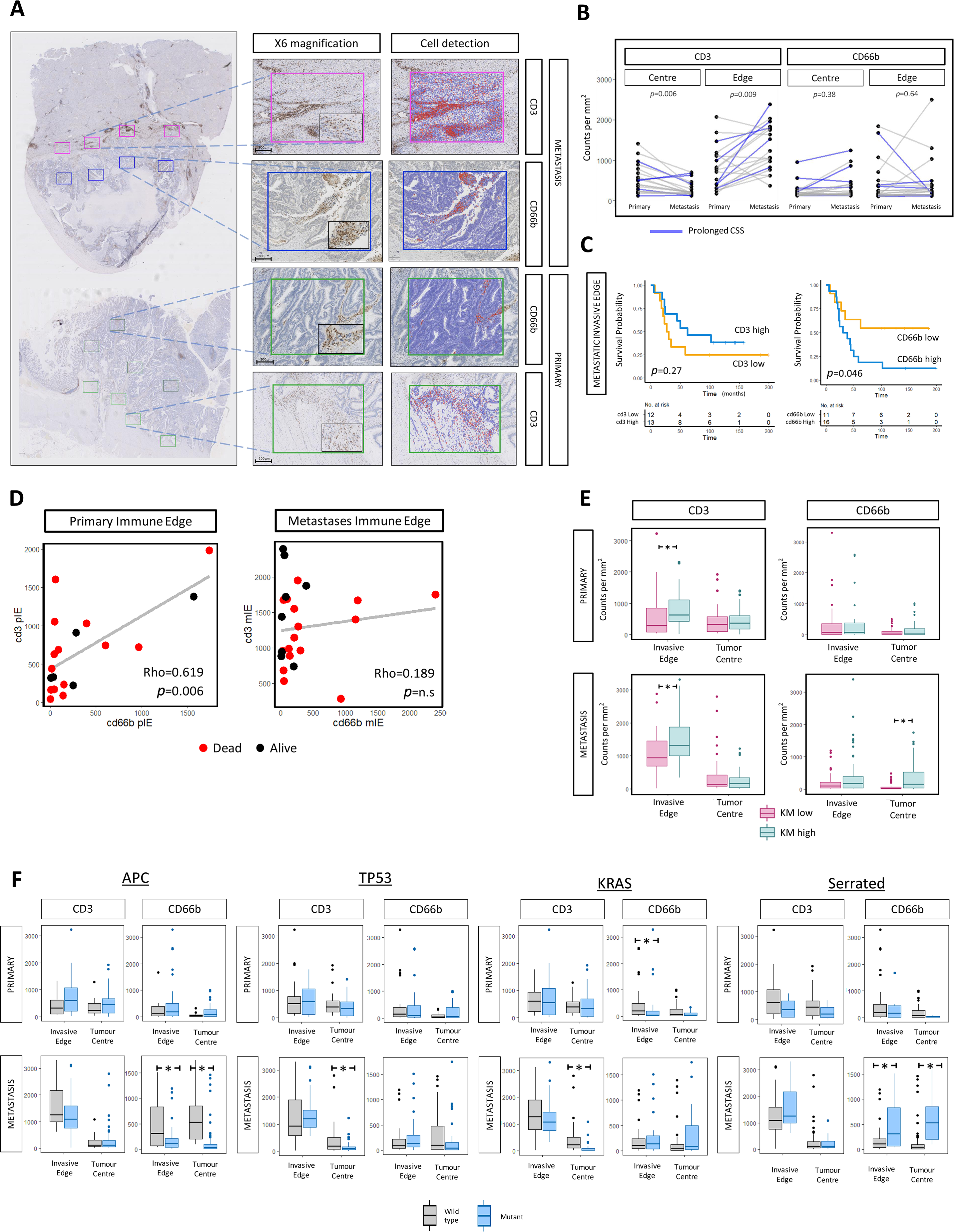
Immunohistochemistry characterisation of matched primary CRC and CRLM with integration of morphological and mutational features. A: Representative images of CD3 and CD66b immunohistochemical staining (Patient B). Whole section demonstrated at 0.5x magnification, ROIs corresponding to tumor centre (TC) and invasive edge (IE) of primary and CRLM shown at x6 and x10 (black box), scale bar 100µm. B: Intra patient comparison between primary and metastasis of CD3 and CD66b cell counts at tumor centre and invasive edge. P values calculated using Mann Whitney test. C: Kaplan-Meier survival plots (log-rank test, P-values displayed) demonstrating the prognostic impact of CD3 and CD66b cell density at CRLM IE identified by IHC for IE of CRLM. High and low values determined according to median expression. D: Comparison of CD3 and CD66b cell density at i) primary IE ii) CRLM IE iii) CD3 primary IE and metastatic TC. Spearman’s Rho analysis. E: Box plots illustrating the relationship between KM grade and CD3 and CD66b cell counts at the IE and TC of primary CRC and CRLM. P value calculated using Mann Whitney test. F: Box plots showing the relationship between mutational features (*APC,TP53,KRAS*, Serrated) and CD3 and CD66b cell density at the IE and TC of primary CRC and CRLM. P values were calculated using Mann Whitney test. Lesions of serrated origin were defined as *APC* wildtype + BRAF mutation.

Survival analysis was performed according to CD3 and CD66b expression across all lesions and regions. CD66b^high^ at the IE of CRLM was associated with worse prognosis (5-year survival 19% vs 64%, Log-rank, *P*=0.046) (Figure 3C).

Co-abundance analysis of CD3 and CD66b demonstrated that within primary CRC IE, CD3 and CD66b were strongly correlated (rho=0.619, *P*=0.006), however, this was not the case in the CRLM (rho=0.189, *P*=n.s) (Figure 3D).

CD3 and CD66b cell density data were then integrated with immune morphological features. CD3 density was elevated at the IE for both CRC (*P*=0.02) and CRLM (*P*<0.005, Figure 3E) in the KM^high^ grade tumors. CD66b density was elevated in the TC of KM^high^ CRLM (*P*<0.005) suggesting that a primed immune response was concordant with infiltration of neutrophils to the CRLM centre. Immune cell abundance and distribution was then assessed according to mutational status, demonstrating that *KRAS* mutation was associated with reduced CD3 density in the TC of CRLM (*P*<0.005) (Figure 3F) suggesting adaptive immune exclusion. APC mutation associated with reduced CD66b density in the TC and IE of CRLM (*P*<0.005). We categorized tumors with *APC* wildtype and *BRAF* or *KRAS* mutation as serrated, a subtype of CRC that derive from serrated polyps via an alternate pathway usually with absence of APC mutation. We found evidence of elevated CD66b density in the IE and TC of CRLM (*P*<0.005) suggesting predisposition to neutrophil infiltration in metastases of serrated origin (Figure 3F).

### Spatially Resolved Transcriptomic Analysis Demonstrates Marked Intra and Inter - Tumoral Heterogeneity and Biological Insights into Prognostic Subtypes of CRLM

Four matched primary CRC and CRLM were selected for ST analysis, the mIF stained images and ROI are demonstrated in Figure 4A. Two of the matched samples were KM^high^ (Patients A and B) and two were KM^low^ (Patients C and D) at primary and metastatic sites, and all samples were Stroma^low^. The IE of both KM^high^ CRLMs were characterized by an α-SMA stained sIE region encapsulating the PanCK stained mTC with abundant CD45+ cells between the border of the sIE capsule and normal liver parenchyma (Supplemental Figures 2,3). Both KM^high^ CRLM also demonstrate tumor necrosis centrally. In contrast, both KM^low^ CRLM had a sharper transition between epithelial mTC and normal liver parenchyma with limited α-SMA and CD45 staining suggesting a less well-defined IE (Supplemental Figures 4, 5).

**Figure 4:**
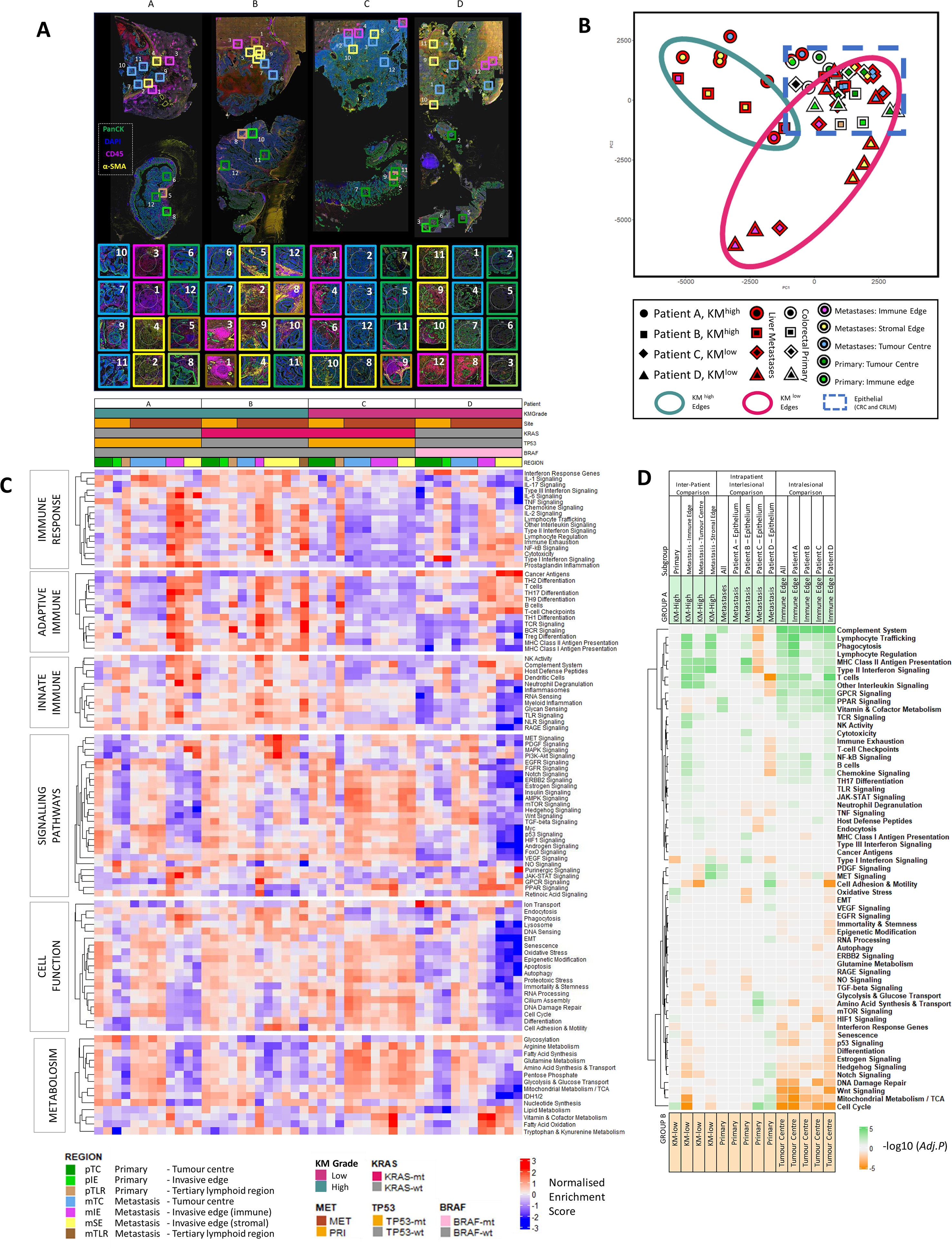
Spatially Resolved Transcriptomic Analysis using Nanostring Cancer Transcriptome Atlas gene sets. A: Representative images of multiplex immunofluorescence (mIF) staining of 4 matched primary CRC and CRLM. DAPI (blue), Pancytokeratin (green), CD45 (pink), a-SMA (yellow). Topographic regions are annotated, each box represents hand selected area of tumour. 8 regions were taken from CRLM and 4 from primary CRC per patient. Patient A: KM^high^ *KRAS*-wt, good prognosis. Patient B: KM^high^, *KRAS*-mt, good prognosis. Patient C: KM^low^, *KRAS*/*TP53* co-mutation, poor prognosis. Patient D: KM-low, *KRAS*-wt, *BRAF*-mt high mutational burden lesion. B: Principal Component Analysis plot of all regions of interest (ROI) selected. The patient from whom the lesion originated is represented by shape. A red border indicates region arises from CRLM and white border represents primary CRC. The topographical region within the lesion is illustrated by the innermost colour of the shape. KM ^high^ metastatic edges are represented by a green circle, KM^low^ metastatic edges are represented by a red circle and the dashed blue line represents epithelial regions of primary CRC and CRLM. C: Heatmap demonstrating single sample GSEA for every ROI, ordered on x-axis by Patient and ROI. Key presented to aid patient identification. Y-axis represents annotated gene sets from Cancer Transcriptome Atlas ordered by and clustered within modules of Immune Response, Adaptive Immune, Innate Immune, Signalling Pathways. Cell Function, Metabolism. Each cell represents the Normalised Enrichment Score scaled by pathway. D: Heatmap demonstrating GSEA providing inter-patient comparison of selected areas between KM^high^ and KM^low^ patients and intra-patient comparison between primary and metastatic sites and intra-lesional comparison between tumour centre and immune edge. Subset of regions filtered prior to GSEA is demonstrated in Subgroup. Subsequent groups compared in GSEA identified as Group A and Group B. Group A annotated at top of diagram and represented by green. Group B annotated at bottom of diagram and represented by orange. Cells of heatmap represent -Log10(Adj.P) for comparison, cell is tinted green if pathway is upregulated in group A and orange if upregulated in group B.

The IO360 bulk transcriptome panel and GeoMx CTA data in matched samples were correlated and grouped by region (Supplemental Figure 9). Gene expression for all GeoMx regions correlated well with the corresponding matched bulk sample (rho 0.612 - 0.906, all P<0.005) however ROIs from primary CRC and the mIE of CRLM correlated most strongly.

A PCA plot of all ROIs demonstrated that most heterogeneity exists in the invasive edges (mIE, sIE) of CRLM, while epithelial regions including TC of primary CRC (pTC) and CRLM (mTC) clustered closely (blue box, Figure 4B). The IEs of KM^high^ and KM^low^ tumors clustered separately for CRLM. However, the mIE from patient C (*KRAS/TP53* co-mutation), clustered more closely with the epithelial group.

ssGSEA was performed across all ROIs employing CTA derived curated gene modules (Immune Response, Adaptive Immune, Innate Immune, Cancer Signalling, Cell Function and Metabolism) demonstrating biological differences both between similarly annotated regions in different patients and between different regions within the same lesion (Figure 4C). To corroborate these observations, DGE and GSEA were performed using inter-patient, intra-patient and intra-lesional sub-group comparisons (Figure 4D). For validation, REACTOME curated gene sets were applied with similar results obtained (Supplemental Figure 10, Supplemental Table 3).

### Intrapatient, Interlesional Heterogeniety Demonstrates Relative Immunosuppression in CRLM in KM ^low^ Patients

Epithelial regions from primary CRC and CRLM within the same patients were compared. While there were no significantly dysregulated gene pathways between primary and CRLM for patient A (KM^high^), in patient B (KM^high^) upregulation of immune signalling in the CRLM included MHC Class II Antigen Presentation (NES=-2.29, *P.Adj*<0.005) and Type II Interferon Signalling (NES=-2.05, *P.Adj*<0.005) (Figure 4D). In KM^low^ patients (Patients C and D), there was relative downregulation of immune related pathways in the CRLM including Lymphocyte Trafficking (NES=1.91, *P.Adj*<0.005) and Type II Interferon Signalling (NES=2.11, *P.Adj*<0.005), Chemokine Signalling (NES=1.76, *P.Adj*<0.005) and T-cell Checkpoints (NES=1.78, *P.Adj*=0.01) (Figure 4D).

### Good Prognosis High Immune Subgroups Are Defined Transcriptomically by Immune Signalling Pathways at Metastatic Invasive Edges

The KM^high^ CRLMs (Patient A and B) demonstrated enrichment of Adaptive Immune and Immune Response CTA gene set modules most profoundly at the mIE with the following altered: B-cell signalling (NES=-1.9, *P.Adj<*0.005); T-cell signalling (NES=-2.05, *P.Adj<*0.005); MHC Class II Antigen Presentation (NES=-2.09, *P.Adj*<0.005); Type II Interferon signalling (NES=-2.05, P.Adj<0.005); Lymphocyte Regulation (NES=-1.81, *P.Adj*<0.005); Lymphocyte Trafficking (NES=-2.06, *P.Adj*<0.005) (Figure 4D). In contrast, patient C (KM^low^ *KRAS*/*TP53* co-mutation) was immune deplete across all regions of the CRLM according to transcriptomic assessment (Figure 4C). While immune signalling was ubiquitous across most immune pathways in the mIE of KM^high^ lesions, in Patient D (KM^low^, *BRAF* mutation) there was reduced expression of Type II and Type III Interferon signalling and MHC Class I and II Antigen Presentation (Figure 4C) however with high Complement System expression at the mIE (NES=-2.68, *P.Adj*<0.005). The KM^low^ CRLMs demonstrate enrichment of atypical immune pathways according to ssGSEA analysis with Patient C having upregulation of RAGE signalling (Innate Immune gene module) with Cancer Antigens and TH9 differentiation (Adaptive Immune gene module) upregulated in Patient D (Figure 4C).

### Poor Prognostic KRAS, TP53 Co-mutated Patient Demonstrates Upregulation of NOTCH and TGF-B Signalling in the Metastatic Tumor Centre

Gene sets within the cancer signalling pathways module were then analysed identifying 3 clusters (Figure 4C). In the first cluster, MAPK, MET, PDGF and PI3K expression appear enriched at the IE of KM^high^ lesions with MET (NES-2.27, *P.Adj*<0.005) and PDGF (NES=-2.14, *P.Adj*=0.01) most significantly expressed at the mSE (Figure 4D). The second cluster was enriched for VEGF, WNT, NOTCH and Hedgehog signalling throughout regions of Patient C and the mTC of Patient D. Comparing the mTC of KM^high^ and KM^low^ lesions, NOTCH signalling (NES=1.72, *P.Adj*=0.022) and TGF-B signalling (NES=1.70, *P.Adj*=0.027) were most significantly upregulated in low immune lesions (Figure 4D). The third cluster included Purinergic and JAK-STAT gene sets which were most highly expressed in mIE and mSE of *KRAS* wild type lesions (Figure 4C).

### KM^low^ CRLM show greater mitotic and metabolic activity than KM^high^ lesions

Next, pathways associated with cellular function and metabolism were analysed demonstrat ing that KM^low^ lesions, particularly Patient C (*KRAS/TP53* co-mutation), were mitotically and metabolically active, particularly at the mTC (Figure 4C). Comparing the mTC of KM^high^ and KM^low^ patients, Cell Adhesion & Motility was significantly upregulated in KM^low^ CRLM (NES=2.11, *P.Adj*<0.005). At the mIE, Cell Cycle (NES=2.21, *P.Adj*<0.005) and Mitochondrial Metabolism (NES=2.36, *P.Adj*<0.005) were upregulated in KM^low^ lesions (Figure 4D).

### Expression of Bulk Transcriptomic Signatures in Spatial Transcriptomic Datasets

The expression of established bulk transcriptomic signatures in this spatial transcriptomic dataset was explored. To date the most comprehensive characterisation of CRLM using bulk transcriptomic techniques was performed by Pitroda *et al* (21) who described 3 CRLM mRNA signatures which were integrated with clinicopathological traits and classified as; 1 - canonical, 2 - immune and 3 - stromal. In this dataset, the genes from all 3 signatures which overlapped with the CTA-panel were markedly over-expressed in the IE of CRLM compared with the TC further demonstrating the capability of ST to demonstrate intra-tumoral heterogeneity and the potential stromal contamination of bulk signatures (Supplemental Figure 11A).

### Spatial Deconvolution of Tumor Regions Demonstrates Distinctive Regional Immune Cell Populations Between KM and Mutational Subgroups

To further interrogate the topographic cellular differences between clinically relevant sub- groups, spatial deconvolution was performed to estimate relative immune cell abundances within ROIs using the SpatialDecon tool (36). Selected validation of the transcriptomic deconvolution analysis was achieved by comparison with chromogenic IHC density data from serial tissue sections (Figure 5A). For CD3 and CD66b we demonstrated that ST immune cell abundance matched closely the IHC cell count (Bland Altman plot, Figure 5B), (rho=0.97 *P*<0.005, Figure 5C). We subsequently illustrated that immune cell populations have distinct spatial distributions with marked immune cell heterogeneity noted between matched primary CRC and CRLM, tumour ROIs and immune subgroups (Figure 6A).

**Figure 5:**
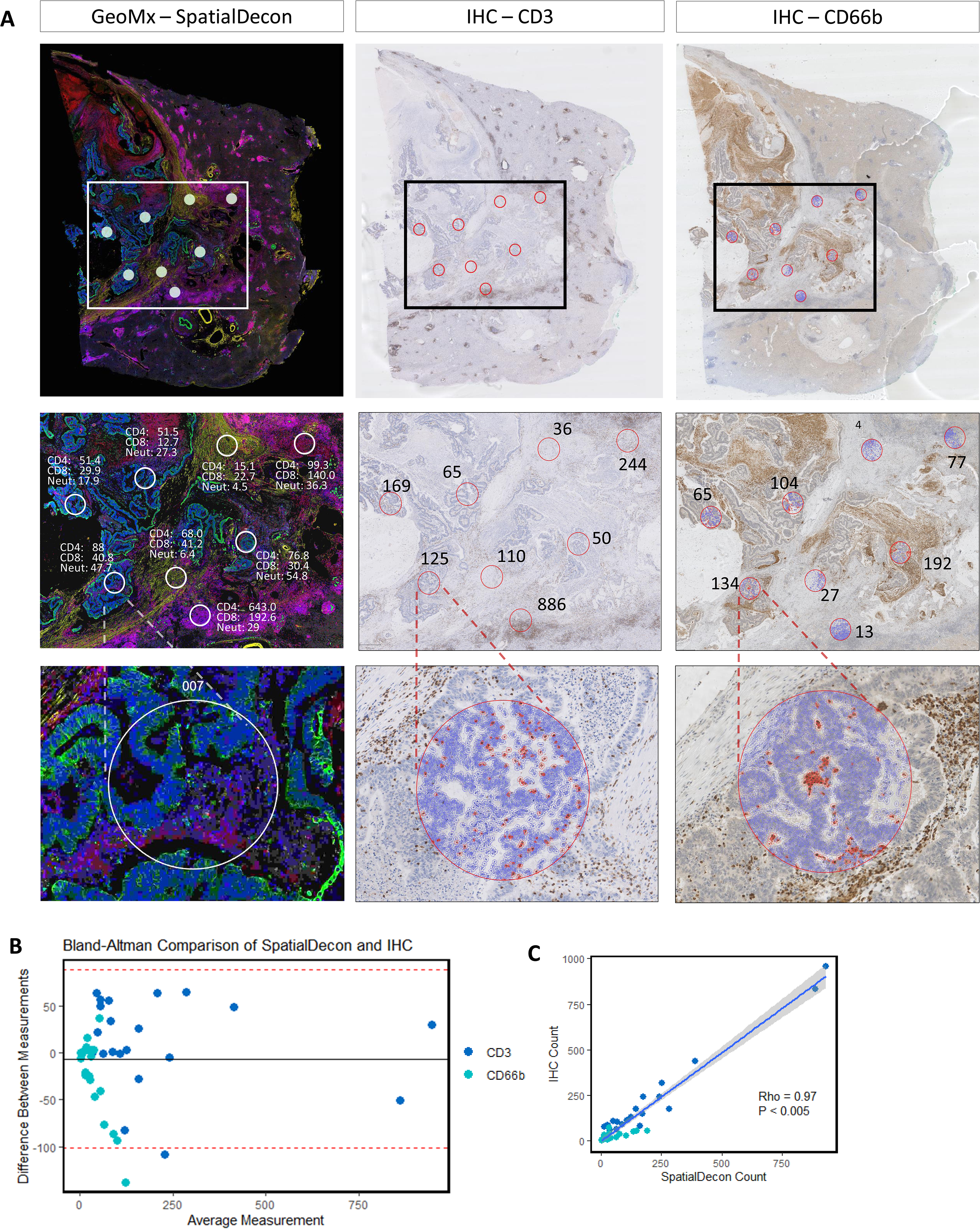
Immune Cell Spatial Deconvolution. A: Representative images from 1 CRLM (Patient A, Figure 4A) showing cell detection from the Qupath package used on a CD3 and CD66b IHC stained liver metastasis to count number of CD3 and CD66b positive cells from 21 regions from a total of 3 CRLM. This count was compared with the SpatialDecon derived count which uses the transcriptomic data from the corresponding ROI in the GeoMx mIF stained matched sample. (See Supplemental Figure 5). B: Bland Altman plot comparing transcriptome SpatialDecon derived cell count versus the IHC derived cell count C: Correlation plot comparing transcriptome SpatialDecon derived cell count versus the IHC derived cell count

**Figure 6:**
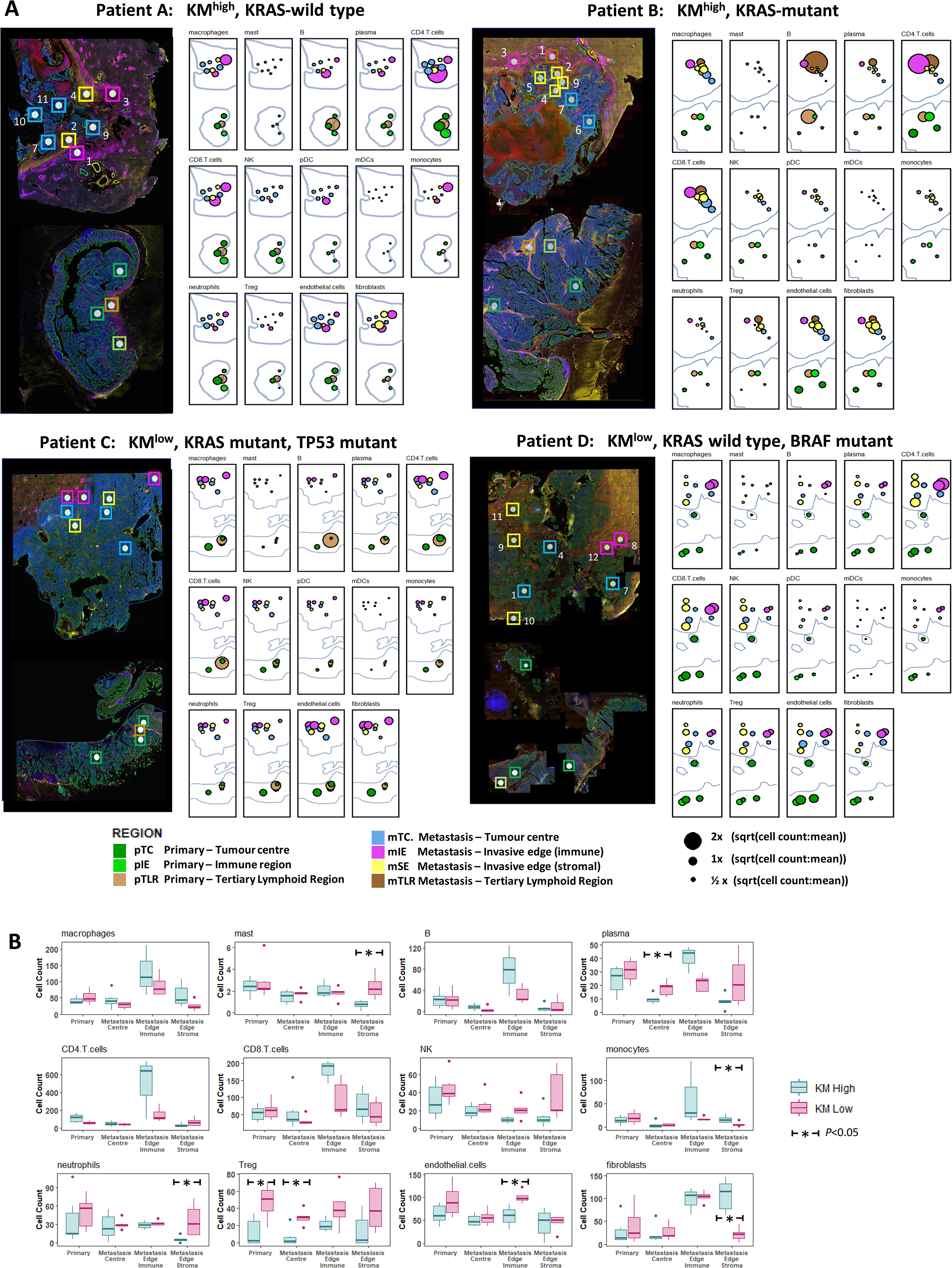
Topographic Immune Cell deconvolution primary CRC and CRLM. A: Images (from Figure 4A) of primary CRC (bottom) and CRLM (top) with 48 ROIs superimposed. Abundance estimates as determined from transcriptome by SpatialDecon for 14 cell populations illustrated for each ROI with colour coding detailing the annotated tumour region. Radius is proportional to the estimated cell counts within the ROI. The immune cell count per region was extracted and the square root of the ratio to the mean immune cell count per region (41.37) of all immune cells was calculated and displayed. The square root of the ratio was calculated to minimise the skew caused by variance of highly expressed cell types. B: Box plots demonstrating the median and interquartile range for each cell type analysed organised by cell type and topographic region and grouped by KM grade. All ROI taken from primary CRC except TLR are grouped as *Primary*. Dendritic cells were removed due to insignificant counts. Mann-Whitney test used to assess for statistical significance. * *P*<0.05

The mIE demonstrated higher abundance of adaptive immune cells compared with mTC (CD4: mean 280.0 vs 46.8, *P*<0.005; CD8: mean 126.8 vs 43.4, *P*<0.005; B-cells: mean 48.5 v 5.8, *P*<0.005) (Figure 6B) (Supplemental Table 4). According to KM status, KM^high^ lesions had higher CD4 (Mean 498.5 vs 150.3, *P*=0.22) and B cells (Mean 77.8 vs 30.9, *P*=0.23) but only CD8 (Mean 180.0 vs 93.5, *P*=0.035) was significantly elevated compared with KM^low^ (Figure 6B). While, regulatory T cells (Treg) density was uniform across topographic tumor regions, according to immunological subtype, KM^low^ tumors demonstrated elevated density compared to KM^high^ tumors (Mean 12.2 vs 40.0, *P*<0.005) across all topographic regions (Figure 6B).

Macrophages demonstrated a topographic distribution similar to CD4 and CD8 cells with higher abundance in mIE regions compared with mTC (mean 103.0 vs 36.1, *P*=0.008) (Figure 6B). However, macrophage density did not differ according to KM status across mIE regions (KM^high^ 127.8 vs KM^low^ 88.1, *P*=0.47). Neutrophils and Natural Killer (NK) cells demonstrated similar relative topographic patterns, with uniform abundance across ROIs.

There were significantly higher levels in KM^low^ tumors (neutrophils: mean 37.8 vs 23.7, *P*<0.005; NK cells: mean 33.1 vs 19.5, *P*<0.005) with the greatest difference observed in the mSE (Neutrophils: mean 36.9 vs 5.74, *P*=0.05; NK cells: mean 37.0 vs 12.7, *P*=0.12) (Supplemental Table 4). The invasive edges of KM^high^ and KM^low^ CRLM also differed with regards to the fibroblast populations with the former characterized by high mSE fibroblast density (Mean 107.9 vs 21.2, *P*<0.005).

The small number of TLRs precluded statistical comparison however we noted the dominant cell types in all TLRs were adaptive immune cells, particularly B-cells, with comparatively higher abundance of macrophages, monocytes and Treg cells in TLR in primary CRC of Patient C (Supplemental Figure 12).

## DISCUSSION

Interest in a personalized oncology approach to the management of CRC incorporating prognostic and predictive biomarkers continues to build, however, heterogeneity of the tumoral immune response may initiate tumor evolution and impede individualized management algorithms. The application of spatial transcriptomic strategies to comprehensively atlas separate compartments including epithelial and microenvironments not only at multiple topographical regions but also temporally through analysis of metastases has potential to deepen our understanding of tumor immune interactions. Furthermore, it may result in an armamentarium of biomarkers to guide management and target therapeutics.

Our analysis of primary CRC and synchronous CRLM has revealed that KM^High^ grade defines a group of patients, regardless of genomic background, with a favourable prognosis following resection of advanced metastatic disease. We therefore demonstrate that in addition to density, location of immune cells is critical to outcome prediction in these patients. The tumor immune geography is not accurately captured by approaches that fail to incorporate the histology or topographic regions of the tumor, including flow cytometry, bulk RNA sequencing and scRNA-Seq. We have demonstrated the fidelity of spatial transcriptomic analysis in this context, enabling accurate and discriminating characterisation of the transcriptome of invasive margins of these tumors both at the primary and secondary sites.

The varying response to surgical resection of CRLM has driven the need for impactful biomarkers. The Immunoscore is a validated immunological metric for primary non-metastatic CRC however the utility in Stage IV disease has only recently been explored. In characterising the immune “multiverse” within a matched primary CRC and CRLM cohort, Van den Eynde *et al* demonstrated comprehensively, as corroborated in this study, vast intra-metastatic, inter-site and inter-patient heterogeneity and postulate this as a potential mechanism for treatment resistance (38). In characterising the invasive margin and tumor centre, the authors demonstrate topographic insights, with higher whole slide immune infiltration in smaller metastases and dense hotspots in larger CRLM. Furthermore, it was noted that in patients with multiple metastases, the immune infiltration of the least infiltrated CRLMs determines outcome most accurately. Several adaptive immune cell markers were characterized and their findings of CD3 abundance in the CRLM invasive margin with pockets of CD20 cells distributed throughout CRLMs was corroborated within our current analysis. The findings reported here supplement this work by demonstrating that innate immune cell populations, particularly neutrophils at the invasive edge, may play a prominent immunosuppressive role in CRLM. The prognostic role of the Immunoscore obtained from the primary lesion in patients with metastatic CRC was not investigated in the work by Van den Eynde et al, however previous data suggest limited benefit (39). This contrasts with the KM score which we have demonstrated remains prognostic regardless of the evaluation within the primary or metastases. To increase the understanding of the tumor immune interface this study focused spatially resolved transcriptional analysis at the pivotal tumour invasive edge which the Immunoscore targets.

Two prognostic subtypes of CRLM have been identified according to broad morpholo gical and biological characterisation. Patients with CRLM experiencing a prolonged survival (KM^high^) following resection are characterized by a fibroblast-rich stromal capsule enriched for PDGF and MET signalling, surrounded by abundant T-lymphocytes with enrichment of MHC Class II Antigen Presentation and Type 2 Interferon Signalling. In contrast, CRLM in patients with a poor prognosis (KM^low^) were characterized by epithelial regions infiltrated by Treg, poorly defined invasive edges infiltrated by neutrophils and sparse adaptive immune cells in combination with overwhelming downregulation of adaptive immune signalling pathways. Spatial transcriptomics further demonstrated distinct differences in the metastatic immune edge of two patients with poor prognosis thus illustrating inter-lesional topographical heterogeneity. In one patient with *KRAS, TP53* co-mutation we observed a devoid immune landscape at the invasive edge. This patient, uniquely in this dataset, demonstrated upregulation of RAGE signalling in the CRLM. A recent in vitro cell-culture study demonstrated RAGE-mediated chemotaxis of immunosuppressive myeloid derived suppressor cells which may offer one possible explanation for our findings in this CRLM (40). In contrast a patient with hypermutated genome and BRAF mutation demonstrated down-regulation of specific immune pathways including Type-II interferon and MHC-Class II antigen presentation highlighting their pivotal importance. Intriguingly, ST demonstrated expression at the invasive edge of this CRLM, upregulation of cancer/testis antigens, a group of antigens expressed only on germ cells or tumours (41), demonstrating the potential for novel discovery and potential patient-specific targets uncovered by a ST strategy.

The findings in this study corroborate previous definitions of three prognostic transcriptomic subtypes of CRLM, one of which was a good prognostic high-immune subtype with demonstrable interferon related pathways (15). Through integrative genomic analysis, we have shown that a TP53/KRAS co-mutated lesion demonstrated profound adaptive immunosuppression, enrichment of NOTCH signalling and TGF-β signalling pathways in the epithelial centre of metastases. This builds upon our recent discovery from a TP53/KRAS co-mutated genetically engineered mouse model (GEMM) in which epithelial NOTCH expression drove aggressive metastatic progression through TGF-β signalling and neutrophil recruitment (37). Interestingly, the analysis of immune cell populations suggests a propensity for neutrophil recruitment to BRAF/KRAS mutant CRLM in the absence of APC mutations, corroborating observations from serrated CRC murine models in human patients (37). Importantly, this GEMM was found to respond favourably to inhibition of TGF-β signalling pathways, limiting metastatic progression. This highlights the potential of a spatial transcriptomic strategy to disentangle the complexity of TME resulting in identification of clinically relevant tumor subgroups vulnerable to immune focussed therapeutic options in the future.

The current study suggests that in the metastatic context, mutational landscape appears to impact outcome less than the host immune response to the CRLM. Through transcriptomic immune cell deconvolution, a marked heterogeneity in immune cell populations was uncovered according to prognostic sub-groups both temporally and spatially throughout a patient’s burden of disease. For CRLM with transcriptomic evidence of immunosuppression, Treg cells appear paramount, as we observed a high density located in epithelial regions of primary and CRLM. Schürch et al recently employed hi-plex CODEX deep-phenotyping to extensively map the immune landscape of primary colorectal cancer highlighting similar importance of immunosuppressive Treg cells, particularly in cellular neighbourhoods where antigen presentation occurs, with a negative effect on outcome (42).

In primary CRC, high stromal content may be indicative of an evolving metastatic process, underpinned by epithelial mesenchymal transition, and associated with poor outcome (43, 44). However, in the current study TSP assessment conferred no prognostic influence when present in the metastatic setting. An affiliate group recently demonstrated that extensive stroma in the primary tumour confounded bulk transcriptomic analysis approaches (17).

They propose that transcriptomic data from targeted areas of the tumor may be an effective strategy to filter out stromal “noise” and more readily obtain pertinent biological data, overcoming the stromal effect also demonstrated in our data in bulk RNA sequencing analysis.

### Limitations

These data were generated from a single-centre, single-surgeon experience in a predominantly treatment naïve cohort limited to synchronously resected CRC primary and CRLM. The power of this cohort is derived from linkage of the primary and secondary site of disease, clearly revealing the prognostic importance of the Klintrup Makinen grade in this context. We acknowledge these pilot data interrogated only a small selection of samples on the GeoMx platform, however, we believe that even in a limited sample size, this technology has demonstrated biological insight which we will now explore with greater power in larger cohorts of CRLM. Whilst the bespoke region selection offered by GeoMx offers flexibility and configurability, the possibility of region selection bias introduced by the user and subsequent lost regions of biological importance is a possibility. Alternative ST platforms exist including the VISIUM (10X Genomics) platform which in contrast provides a more comprehensive topography of a single larger area (6.5mm^2^) per section through sequencing of 5000 barcoded dots (45). In this study the CTA has offered insight into differences in immune and cancer signalling pathways between prognostic groups, however employment of a whole transcriptome pipeline may have facilitated a more powerful discovery approach.

Finally, our region selection strategy did not employ segmentation, and therefore each region had mixed epithelial and stromal components. Our future GeoMx experiments will take advantage of more advanced segmentation and cell detection strategies.

## Conclusion

In conclusion, the invasive edge of CRLM influences outcome in this resected cohort and can be thoroughly characterized using spatial transcriptomic analysis, and in future may allow researchers to fully address the reason for resistance to therapies and recurrence following surgery. This spatial transcriptomic study of a synchronously resected cohort of CRC and CRLM has demonstrated novel insights into the pathways driving tumor progression on an individual patient basis. This study illustrates the potential to leverage spatial transcriptomics performed on the GeoMx DSP to characterize tumor heterogeneity, and identify novel biomarkers associated with clinically relevant subtypes of CRC. This is a highly utilisable platform in formalin fixed paraffin embedded tissues from human and pre-clinical models.

There was high concordance between spatial immune cell deconvolution with IHC protein assessment and future work will seek to integrate deep immuno-phenotyping, transcriptomic output and multi-omic integration, with a view to applying this technology at scale in addition to serial analysis of lesion through interrogation of biopsy specimens.

## AUTHOR CONTRIBUTIONS

Conceptualization, Methodology, Writing – Original Draft: CSW, KAF, JE, CWS, NBJ. Visualisation, Formal Analysis: CSW, KAF.

Software: CSW.

Writing – Review & Editing: all authors.

Investigation – CSW, KAF, HL, AL, AJC, LMS, HCVW, JE, CWS, NBJ

Resources, Data Curation: SC, JH, CN, AVB

Supervision, Project Administration, Funding Acquisition: CSDR, DCM, OJS, PGH, JE, CWS, NBJ.

## ACKNOWLEDGEMENTS

We gratefully acknowledge NHS Research Scotland GGC Biorepository and Glasgow Research Tissue Facility with their role in accessing tissue for this study.

The DSP GeoMx experiment data was generated in collaboration with the Nanostring technology access program team in Seattle.

## DATA AVAILABILITY STATEMENT

The data generated in this study are available upon request from the corresponding author.

## Supplemental Methods

### Immunohistochemistry

Sections were dewaxed in histoclear and rehydrated through a series of graded alcohols. Antigen retrieval was performed using TRIS-EDTA buffer pH8 for CD3, TRIS-EDTA buffer pH9 for CD66b. Sections were heated in respective buffer for 5 mins under pressure and then cooled for 30 mins. Endogenous peroxidase activity was blocked by adding slides to 3% hydrogen peroxide for 20 mins. Sections were washed in running water and then blocked for 1 hour at room temperature using 5% goat serum. Sections were incubated overnight at 4°C in primary antibody diluted in antibody diluent (Dako) at the following concentrations: CD3 (Sigma) 1:4000, CD66b (Novus) 1:4000. After overnight incubation, sections were washed in tris-buffered saline (TBS) and ImmPRESS reagent was applied for 30 mins at room temperature. Sections were washed in TBS and DAB substrate was added for 5-10 mins before washing in running water. Counterstaining was performed using haematoxylin for 5 minutes, rinsed in water, 1-2 seconds in acid-alcohol, rinsed in water, 45 seconds in Scotts tap water substitute. Sections were dehydrated through a series of graded alcohols, passed through histoclear and coverslips were mounted using distrene plasticizer xylene.

### Nanostring nCounter PanCancer IO360 Bulk Transcriptomic Platform

#### RNA extraction from FFPE tissue

RNA was extracted from the samples using AllPrep DNA/RNA FFPE Kit (Qiagen) according to manufacturer’s protocol, using xylene for deparaffinisation. All RNA samples were treated on-column with DNAse I and were eluted in 20ul RNAse-free H2O. RNA quantity was assessed using RNA BR assay and the Qubit^®^ 2.0 fluorometer (Invitrogen, Life Technologies). RNA integrity (RIN) values were determined using Agilent 2100 Bioanalyzer (Agilent Technologies) (maximum RIN = 2.1).

#### Gene Expression Profiling using NanoString Immune Oncology (IO 360) Panel

Total RNA (Average concentration = 35ng/µl) was used for gene expression analysis using the commercially available IO360 panel (nCounter, NanoString) (770 genes). Each 5µl RNA sample was analysed in sets of 11. Panel standards were used in each assay to allow for differences in technical assay performance across runs. Samples and panel standards were spiked with positive control probes ranging from 128fM to 0.125fM in a four-fold dilution series. Positive controls were used to assess the linearity of the assays and used for normalisation. Additionally, eight negative control probes were used for which no RNA target was present. RNA samples were incubated for 16 hours at 65°C in hybridisation buffer containing IO360 CodeSet, including reporter and capture probes in addition to target RNA forming a tripartite hybridisation complex. Hybridized samples were processed using the nCounter Prep Station (High Sensitivity Protocol), in accordance with IO360 panel guidelines by Nanostring. Data acquisition was performed by using the Nanostring’s Digital Analyser (FOV, 555).

#### Gene expression analysis

Raw gene expression count data was normalized in NanoString nSolver 4.0 using 6 positive controls and 8 negative controls to account for background noise and sample variation across several runs performed on the nCounter platform. Background threshold was manually calculated using Mean (negative controls) ±2 standard deviations of negative control probes to remove lowly expressed targets prior to gene expression analysis. Then, the nSolver 4.0 software enabled data normalisation to be performed using the GeNorm Algorithm with integrated housekeeping gene probes.

#### Nanostring GeoMx Digital Spatial Profiling

##### Slide preparation and hybridisation with UV-photocleavable CTA probes

The four matched 5μm-thick FFPE colon cancer and corresponding liver metastasis tissues were co-mounted on Superfrost glass slides (Thermofischer). The slides were baked for 30 mins at 60 °C. The tissues were dewaxed, hydrated and treated with 1μg/ml Proteinase K for 15 minutes. Next, heat-induced epitope retrieval (HIER) was performed on the Leica BOND Autostainer (ER2 for 100°C) for 20 minutes using a pH 9 antigen retrieval buffer (Leica BOND). The slides were immediately stored in 1x PBS. The tissues were then hybridized as per manufacturer’s protocol with a pre-designed panel of antibodies corresponding to 1,825 genes, known as the Cancer Transcriptome Atlas (CTA) (Nanostring). The CTA is a panel of 1825 genes designed to comprehensively characterize immune activity and tumor biology within tumor microenvironments (46). Each tissue was covered with 200μL of hybridization solution and a HybriSlip™ cover and incubated overnight at 37 °C for at least 16 hours. The slides were then dipped in 2x SSC-T and washed twice with a 1:1 ratio of 100% deionized formamide (Ambion) and 4x SSC at 37°C for 25 minutes each. The slides were blocked with Buffer W (Nanostring) before the addition of fluorescently-labelled morphology markers on the tissue to highlight the tissue’s architecture. The GeoMx DSP hosts four channels (FITC/525nm, Cy3/568nm, Texas Red/615nm and Cy5/666nm) for the detection of up to four customisable morphology markers for each tissue (20). One channel is reserved for the nuclear stain (DAPI), which left three free channels. Consistent with the aim of this project, the additional morphology markers were Pan-Cytokeratin (PanCK) to stain the cytokeratin in the epithelium, cluster of differentiation, CD45, for the immune cell populations and α-SMA to identify fibrogenetic collagenous architecture in liver tissue. The slides were then stored at 4°C in SSC before being loaded on the GeoMx DSP instrument for region selection and collection.

##### Probe collection and Library Preparation

Once region selection was complete, collection was initiated using the DSP workstation whereby photo-cleavable oligonucleotide probes in the user-defined regions were exposed to UV-light to cleave the UV-sensitive probes which releases the probe-specific DSP barcodes which are aspirated from selected regions and dispensed. Following collection, the DSP plate containing the probes was dried and rehydrated with DEPC-treated water before each sample was added to the corresponding well of a new 96-well PCR plate containing the GeoMx Seq Code primers and the PCR Master Mix. A thermocycler was user to incubate the PCR plate using manufacturer settings (Supplemental table 1). The products of the PCR reaction were then centrifuged and pooled prior to purification which was performed using AMPure XP system. Once purified, the library was resuspended in Elution Buffer (10mM Tris-HCl with 0.05% Tween-20, pH 8.0 prior to QC using an Agilent Bioanalyser. Having met Nanostring’s recommendation, the experiment proceeded to NGS.

#### Sequencing

The purified library was loaded onto a flow cell and placed on the Illumina NextSeq 550, where the RNA fragments were amplified in a process called cluster generation. Then, during sequencing by synthesis (SBS), fluorescently-labelled complementary nucleotides bind and read the RNA sequence in a two-way fashion – first forward and then reverse – through a method known as paired-end sequencing generating. The GeoMx NGS pipeline hosted in the Illumina BaseSpace platform was used to convert sequenced FASTQ files into DCC files which can be uploaded onto the GeoMx DSP analysis suite. Once loaded, they undergo quality control, filtering, Q3 normalisation and background correction. The counts were then downloaded from the GeoMx instrument and loaded in to RStudio (v1.2.1335) using R build version 4.1.1 for analysis.

## LIST OF ABBREVIATIONS

CRC: Colorectal Cancer
CRLM: Colorectal Cancer Liver Metastases
IHC: Immunohistochemistry
KM: Klintrup Mäkinen
TSP: Tumor Stroma Percentage
scRNAseq: single-cell RNA sequencing
ST: spatial transcriptomics
DSP: digital spatial profiling
TME: tumor microenvironment
MIF: multiplex immuno-fluorescence
FFPE: Formalin-Fixed Paraffin-Embedded
DFS: Disease-free survival
RFS: Recurrence-free survival
OS: Overall survival
CSS: Cancer-specific survival
HR: Hazard ratio
OR: Odds Ratio
CI: Confidence interval
PH: proportional hazard
MDT: multi-disciplinary team
TIL: tumor infiltrating lymphocyte
GSEA: gene set enrichment analysis
ORA: over representation analysis
H&E: hematoxylin and eosin
n.s: not significant
IE: invasive edge
TC: tumor centre
mIE: metastatic invasive edge – immune
sIE: metastatic invasive edge – stromal
mTC: metastatic tumor centre
mTLR: tertiary lymphoid region (metastasis)
pTC: primary CRC tumour centre
pIE: primary CRC invasive edge
pTLR: tertiary lymphoid region (primary)

## Supplementary Figures

**Supplementary Figure 1:**
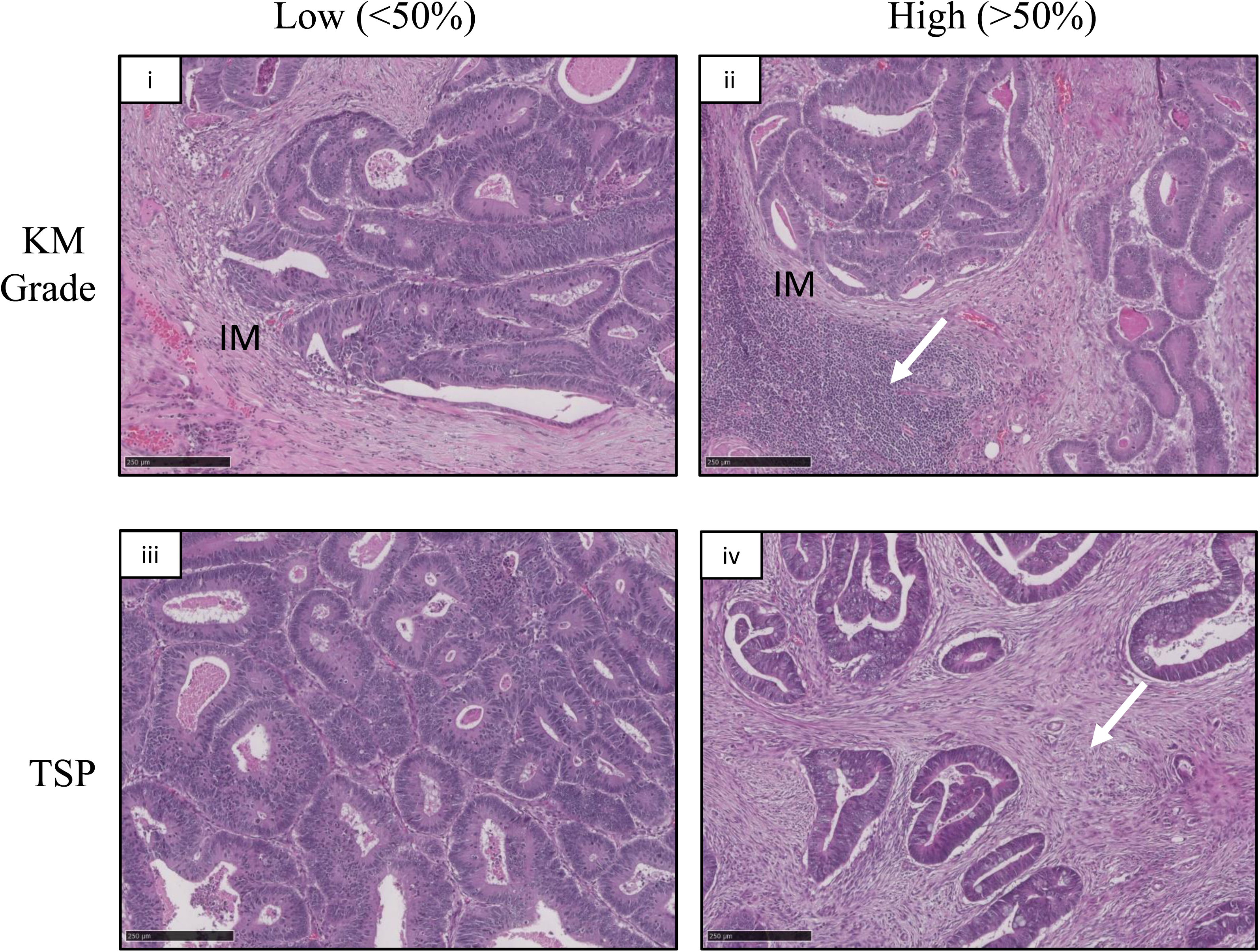
Representative images of H&E sections for scoring of Klintrup-makinen KM grade and Tumor stroma percentage) TSP of primary colorectal cancer. Images taken at x10 magnification (NDP Viewer software), scale bar 250µM. (i) KM^low^ graded tumor (ii) KM^high^ tumor (high immune infiltration arrowed). (iii) low intra-tumor stromal percentage (TSP) (<50%) (iv) high (>50%) TSP tumour (stromal area arrowed).

**Supplemental Figure 2:**
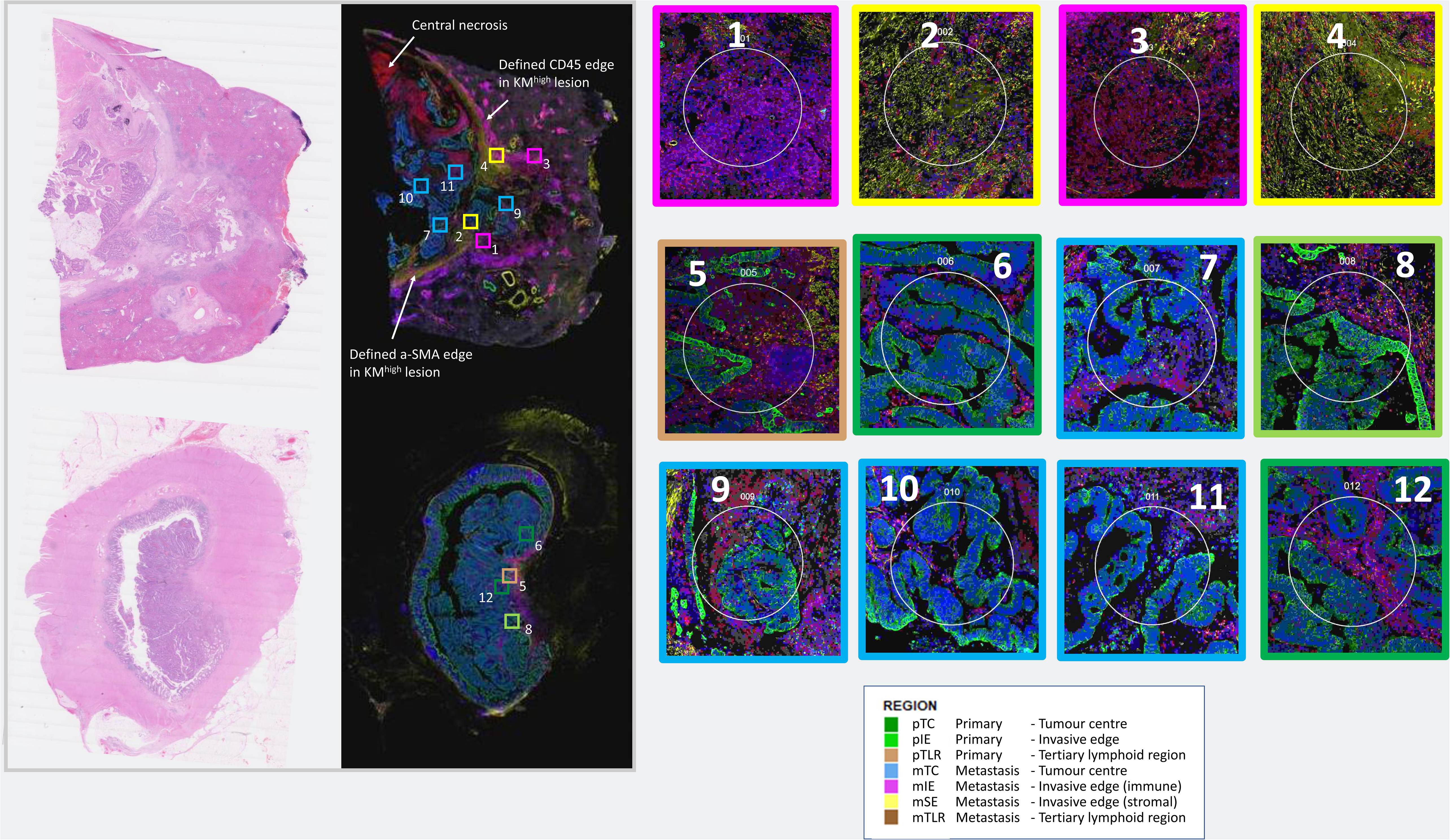
Patient A: KM^high^, KRAS wildtype type. Representative H+E and mIF stained sample with annotated regions colour coded (from Figure 4A, 5A and 6A) alongside magnified image of regions, 4 primary and 8 metastases, from patient A from GeoMx experiment.

**Supplemental Figure 3:**
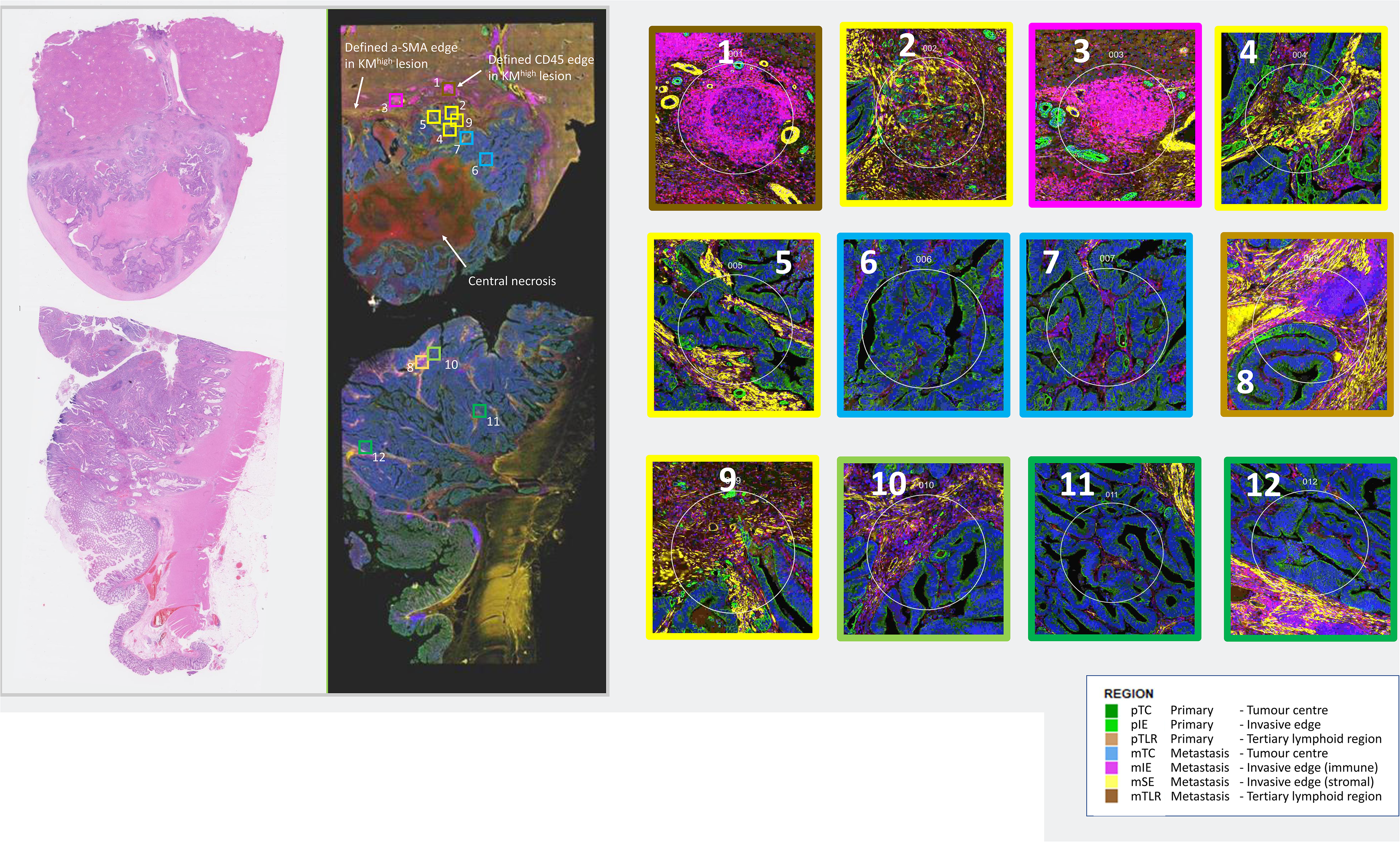
Patient B: KM^high^ immune, KRAS mutation. Representative H+E and mIF stained sample with annotated regions colour coded (from Figure 4A and 6A) alongside magnified image of regions, 4 primary and 8 metastases, from patient B from GeoMx experiment.

**Supplemental Figure 4:**
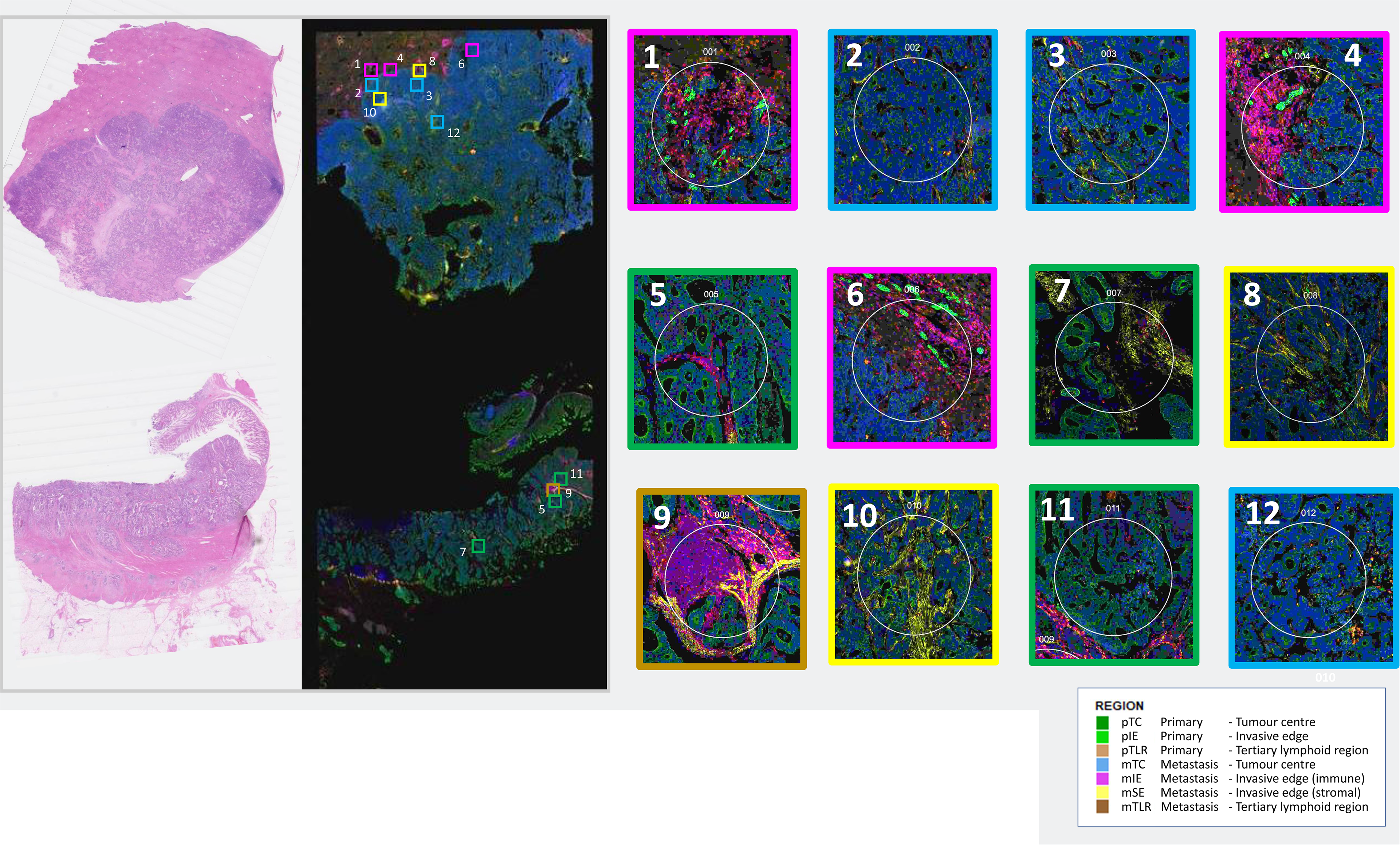
Patient C: KM^low^ immune, KRAS mutation. Representative H+E and mIF stained sample with annotated regions colour coded (from Figure 4A and 6A) alongside magnified image of regions, 4 primary and 8 metastases, from patient C from GeoMx experiment.

**Supplemental Figure 5:**
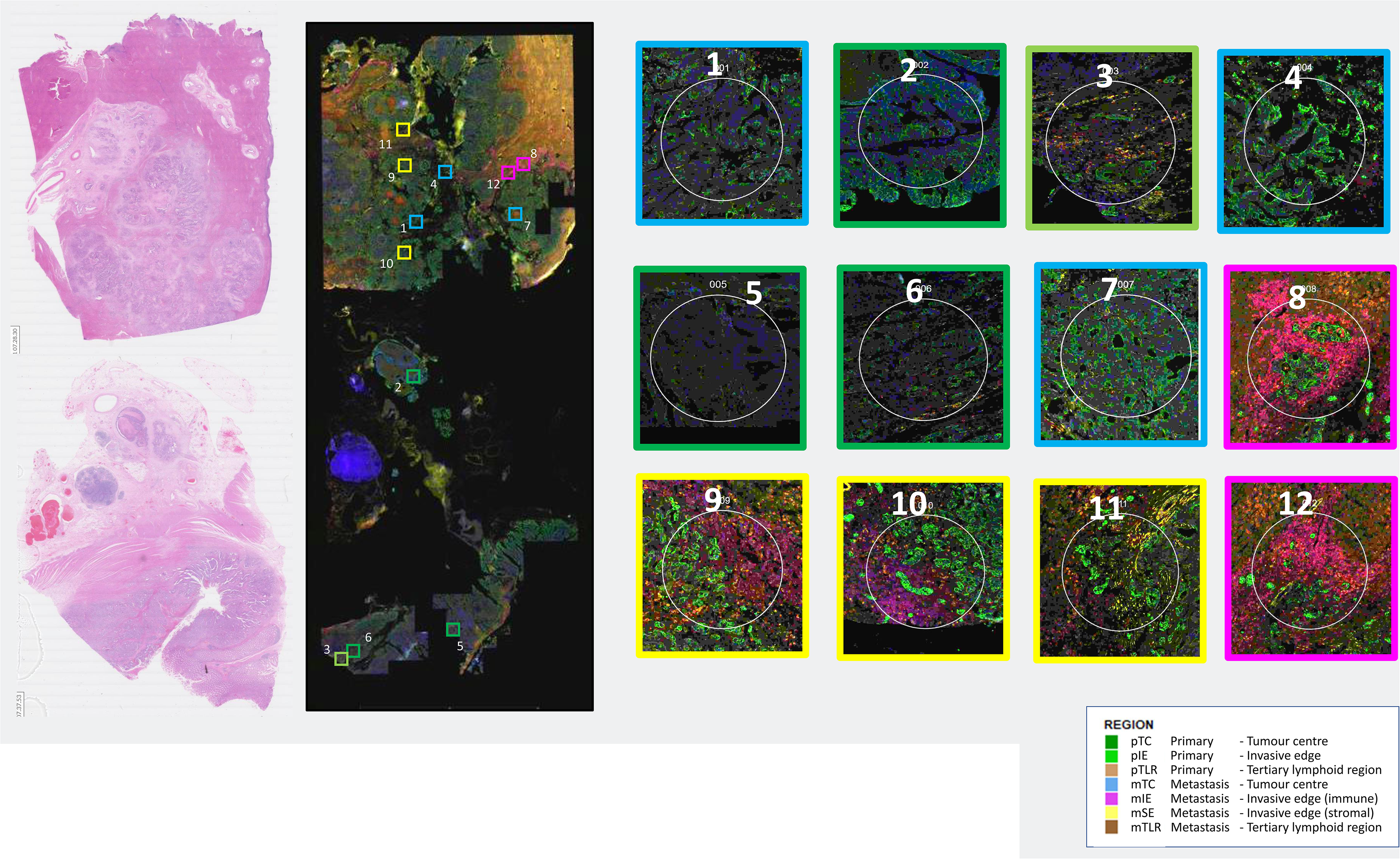
Patient D: KM^low^ immune, KRAS wildtype, BRAF mutant. Representative H+E and mIF stained sample with annotated regions colour coded (from Figure 4A and 6A) alongside magnified image of regions, 4 primary and 8 metastases, from patient D from GeoMx experiment.

**Supplemental Figure 6.**
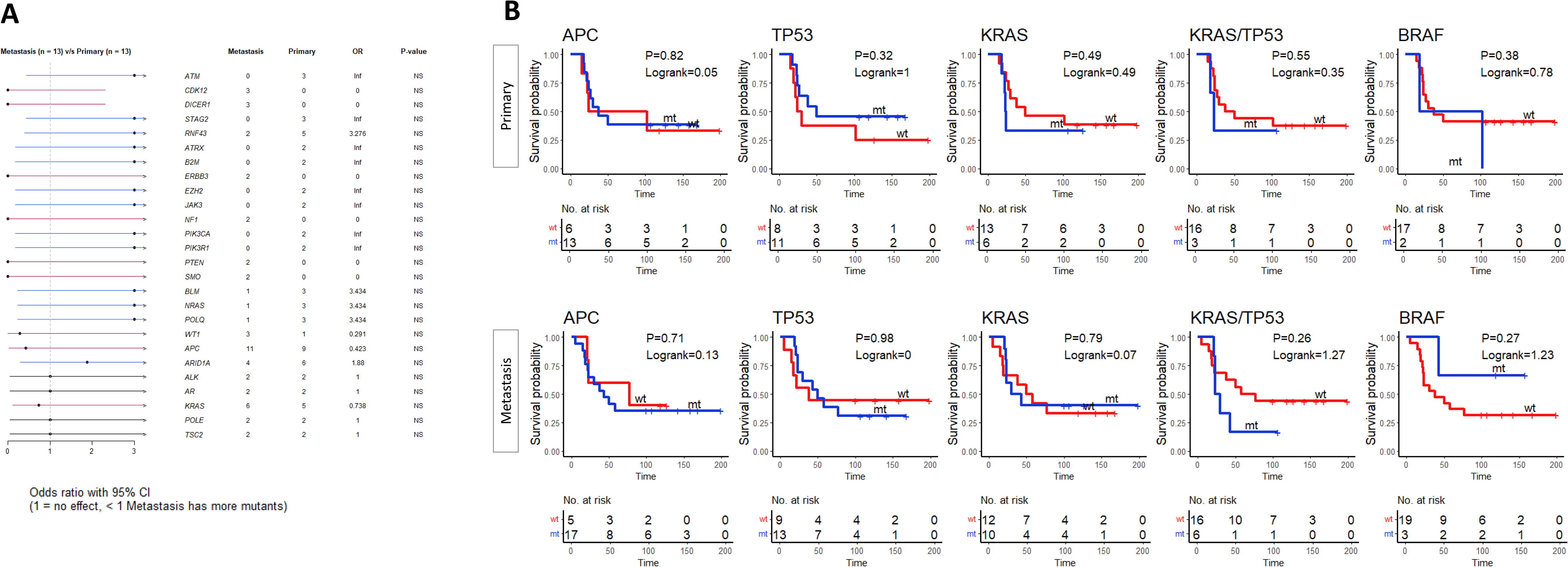
A: Forrest plot demonstrating mutational discordance between primary and secondary (non -matched lesions). B: Kaplan Meier survival plots for cancer specific survival by mutational groups with Log-rank test and P-value displayed.

**Supplemental Figure 7:**
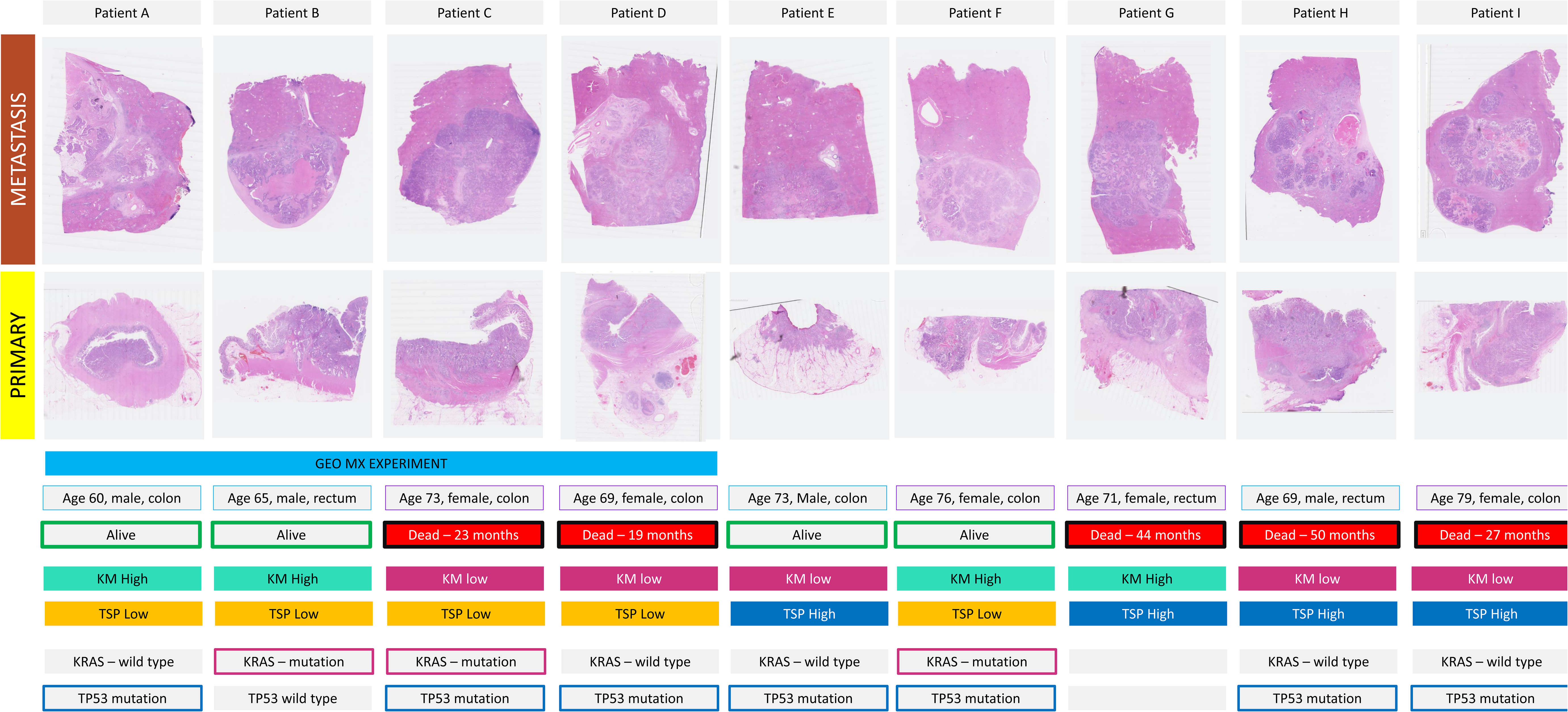
Representative H+E and clinicopathological, mutational and survival details of patients (n=9) undergoing Bulk transcriptomic analysis on the nCounter platform. 9 patients from the cohort (patients A-D in Supplemental Figures 2-4) had bulk transcriptomic analysis on the ncounter platform using the Pancancer IO360 (770 genes) panel with clinicopathological, morphological and mutational characteristics as demonstrated. Representative H+E from the primary and metastatic lesions from the patients are demonstrated. 4 of the patients were selected for analysis on the GeoMx Digital Spatial Profiler. A selection of KM and TSP score were selected for comparison.

**Supplemental Figure 8.**
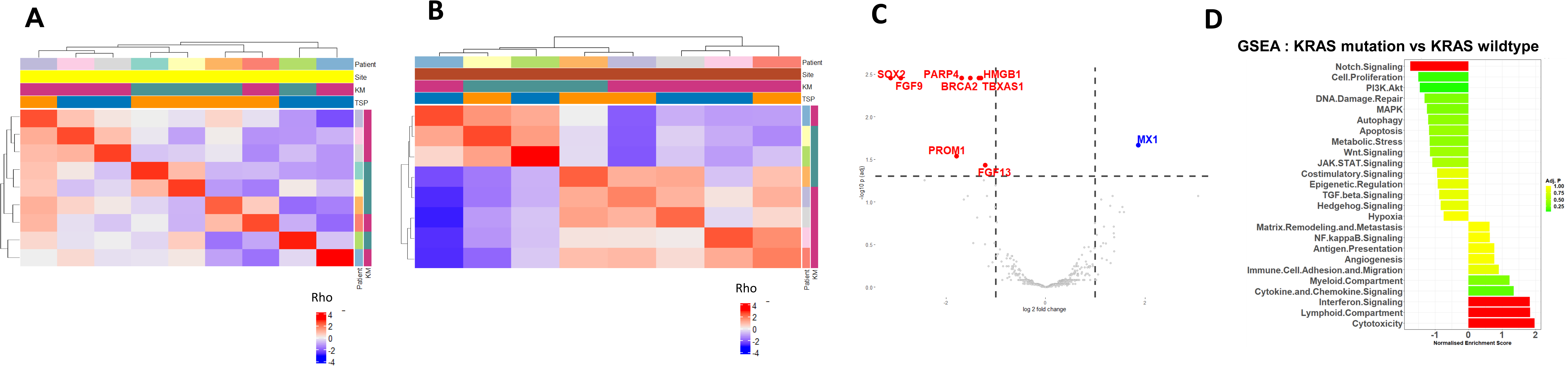
A: Unsupervized analysis using gene expression correlation matrix for Primary lesions only. Patient, site, KM grade, tumor stromal percentage (TSP) are depicted by key. Spearman correlation of all expressed genes performed between each sample sequenced and plotted on the heatmap. k -means clustering of heatmap to demonstrate correlated samples. Red represents strong correlation. Blue represents negative correlation B: Unsupervized analysis using gene expression correlation matrix for Metastatic lesions. Patient, site, KM grade, tumor stromal percentage (TSP) are depicted by key. Spearman correlation of all expressed genes performed between each sample sequenced and plotted on the heatmap. k -means clustering of heatmap to demonstrate correlated samples. Red represents strong correlation. Blue represents negative correlation. C: Volcano plot representing differential gene expression between KRAS-wildtype and KRAS-mutant lesions. X-axis represents Log2Fold Change, Y-axis represents –Log10(Adj.P). D: Bar chart representing Normalized Enrichment Score and Adjusted P-value from Gene Set Enrichment Analysis performed on nCounter gene sets for ranked differentially expressed genes between KRAS-wildtype and KRAS-mutant lesions.

**Supplemental Figure 9:**
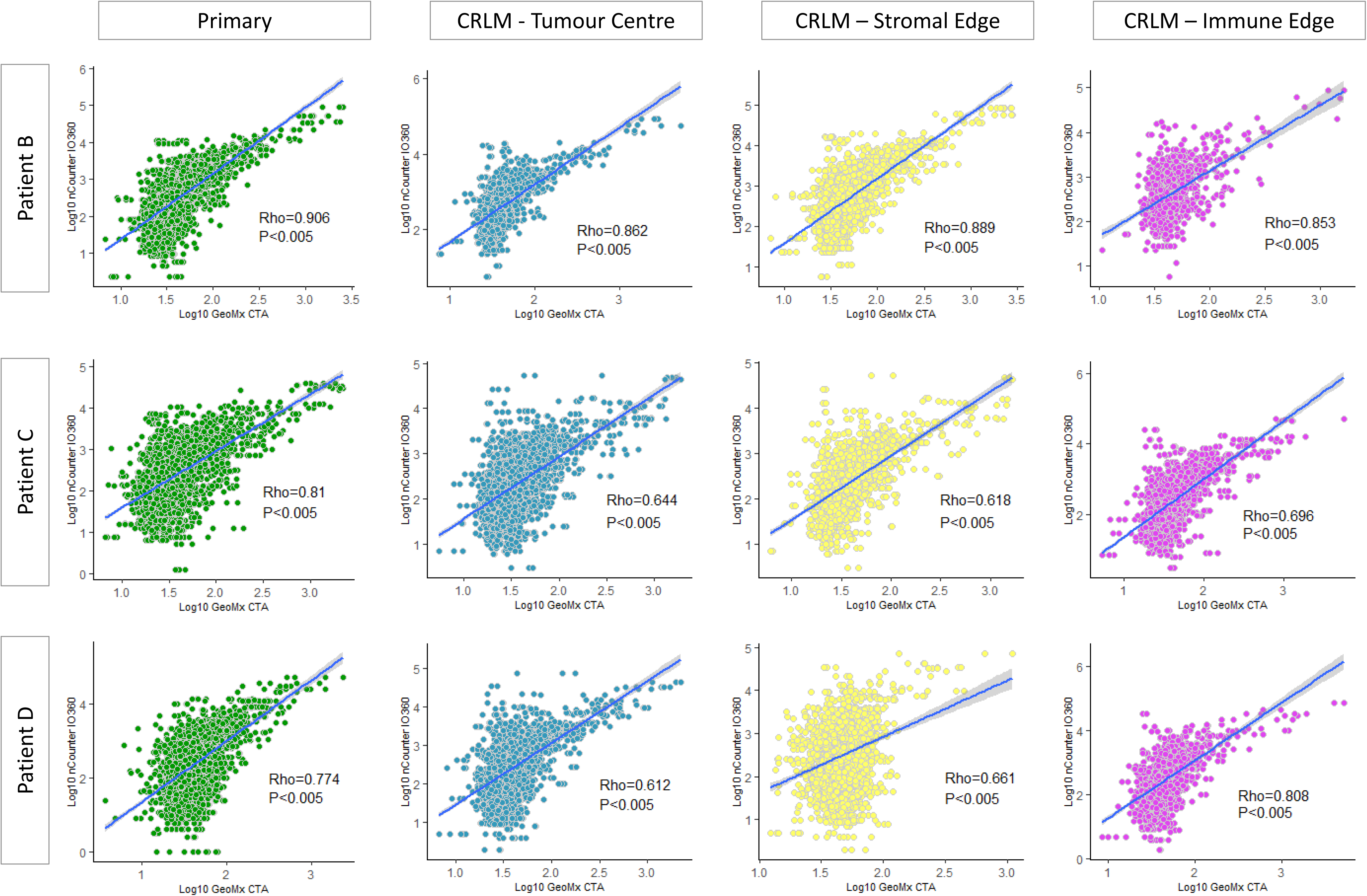
Comparison of nCounter IO360 and GeoMx CTA Gene Expression in Matched Samples. Each dot represents log10 expression of gene on nCounter IO360 (y-axis) and GeoMx (x-axis) organized by region and patient (Patients B-D)

**Supplemental Figure 10:**
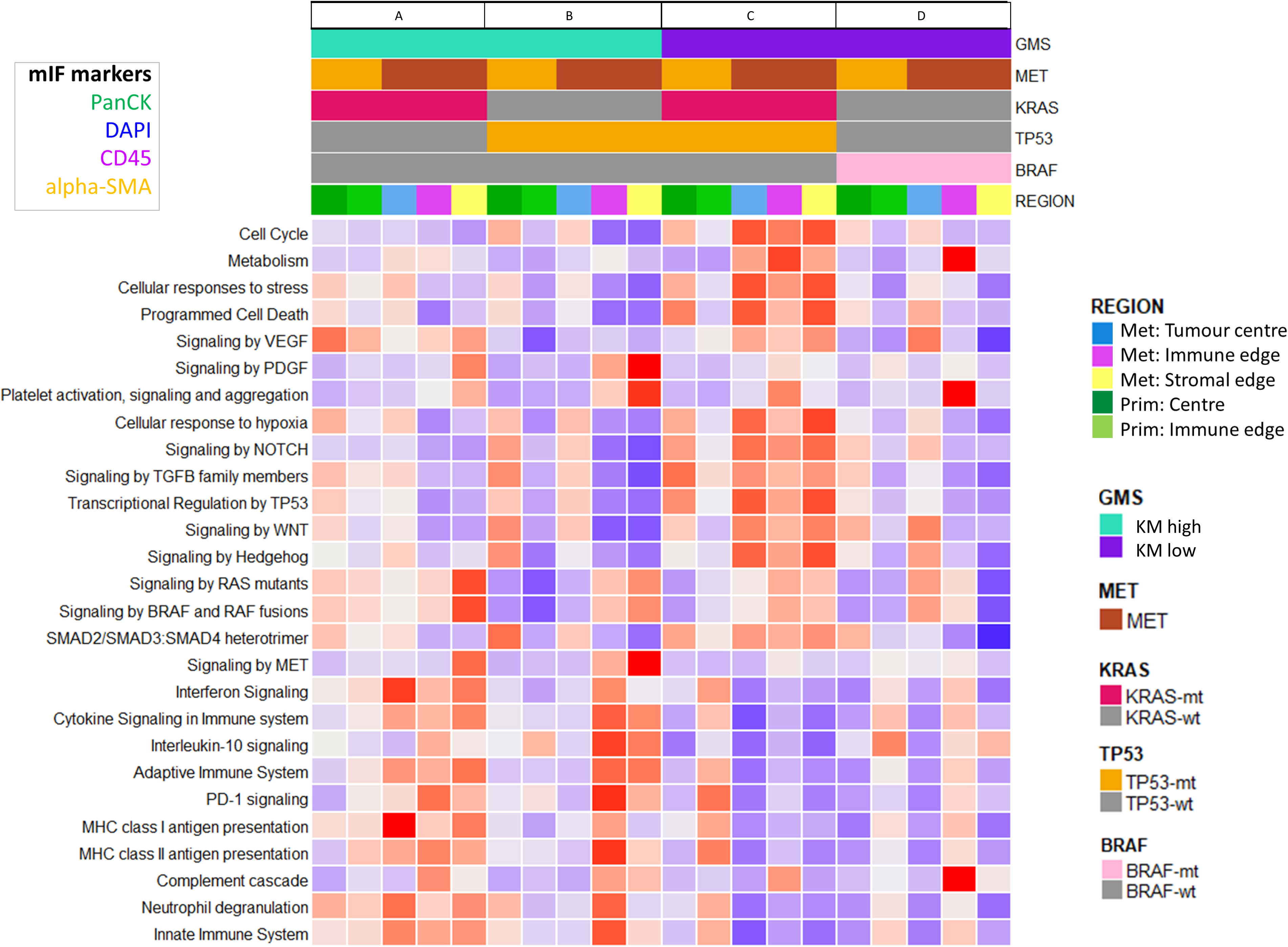
Spatially Resolved Transcriptomic Analysis using REACTOME curated gene sets. Unclustered heatmap demonstrating summarized mean gene expression for each selected REACTOME pathway with annotations as shown. Ordered by patient, lesion (primary, metastasis) and annotated topographical region.

**Supplementary Figure 11.**
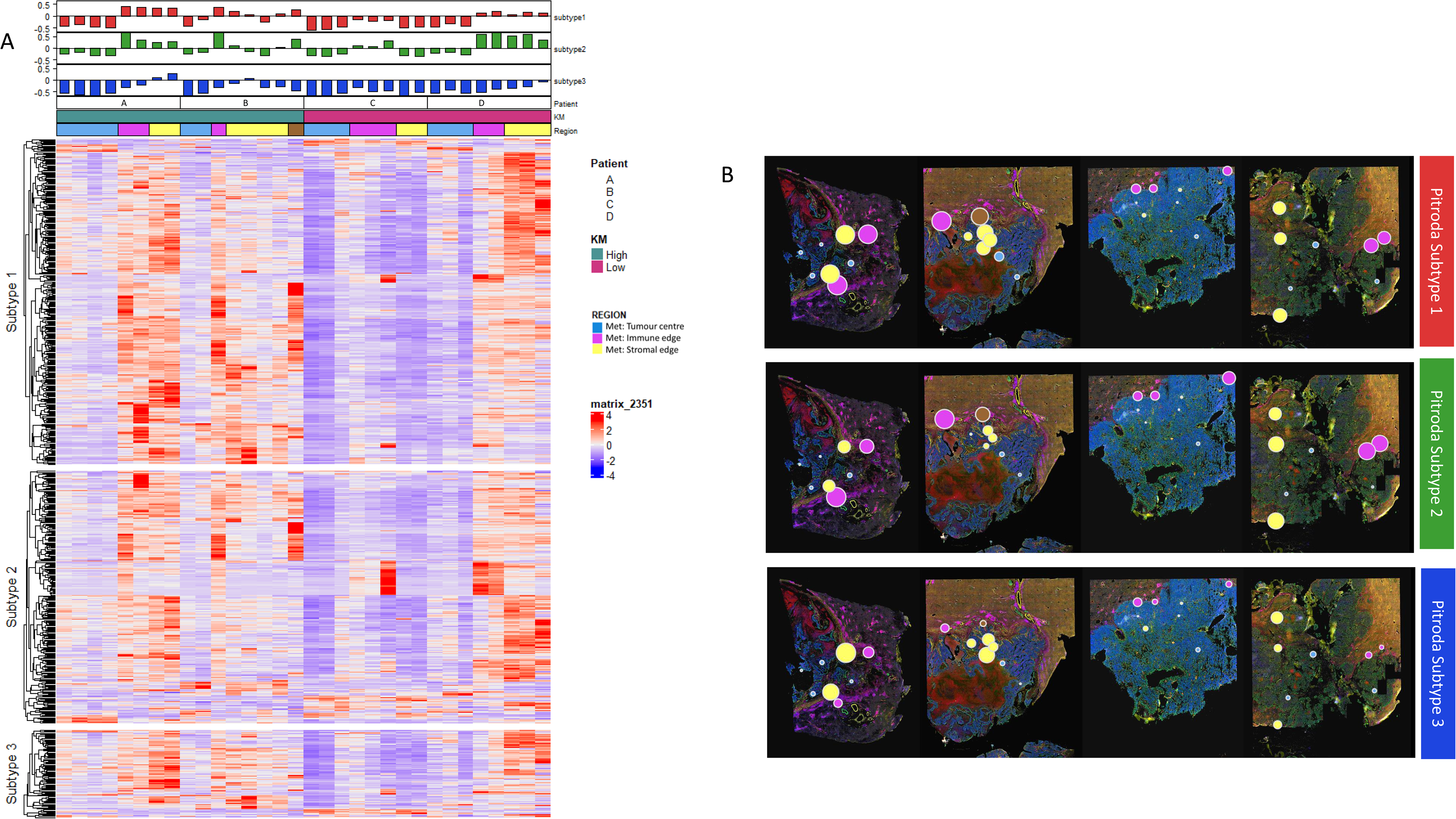
A: Pitroda et al gene sets applied across spatial transcriptomic data set. ssGSEA enrichment score is at the top of the heat map. Heat map comparing expression across CRLM and region. Heat map comparing expression across CRLM and region. Overlap between CTA and Pitroda geneset, Subtype 1: (348 of 1506 genes), Subtype 2: (270 of 1259 genes) and Subtype 3: (94 of 603 genes). B: Geneset Enrichment score for each Pitroda subtype demonstrated by radius of region and topographic location on tumour. Annotated region demonstrated by color (images also shown in Figure 4A and 6A and Supplemental Figures 5-8).

**Supplementary Figure 12:**
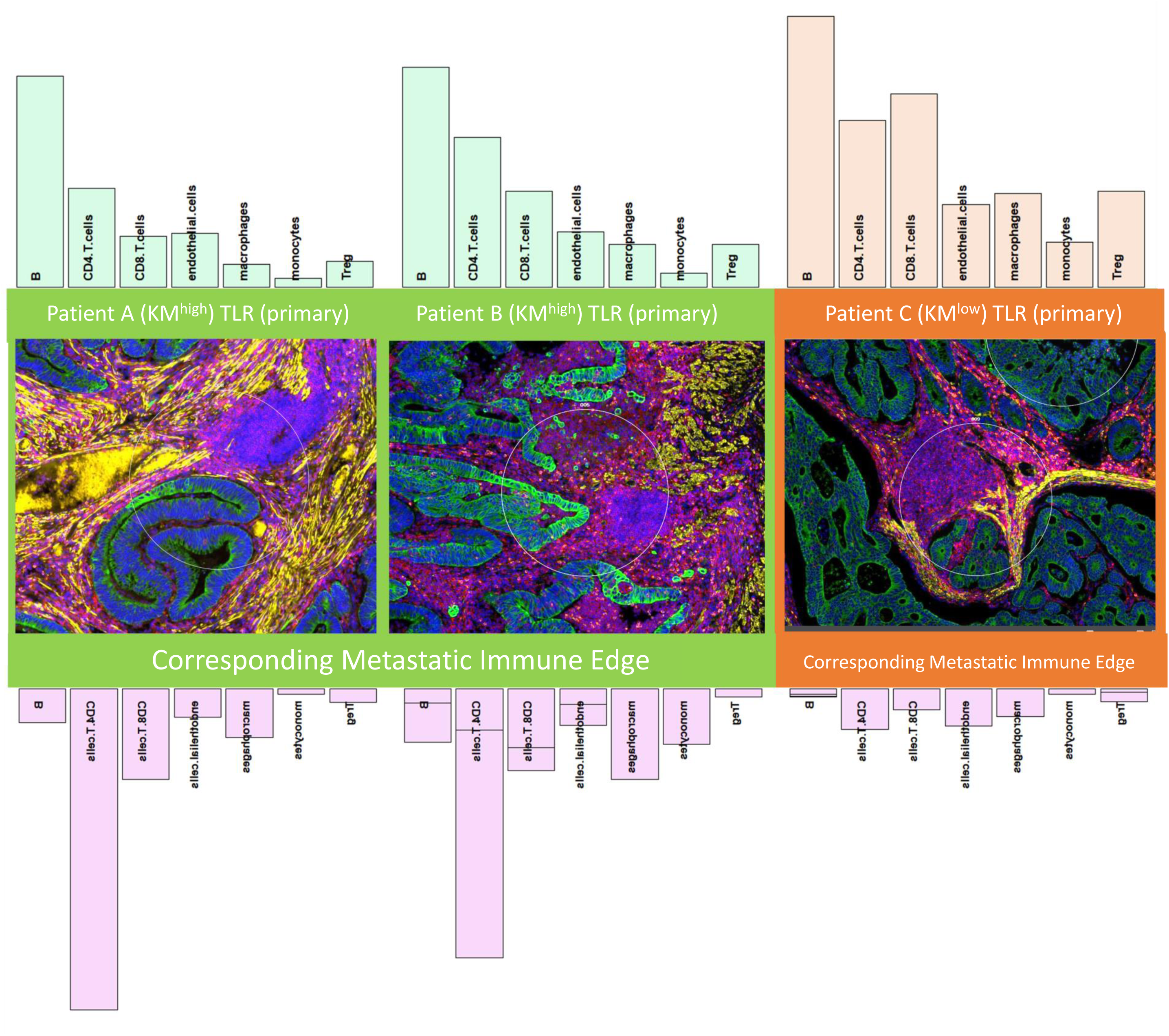
TLR Immune Cell Spatial Deconvolution. The immune cell populations are plotted for Tertiary Lymphoid Regions in Lesions A,B (KM-high) (Patient B, image 8, Supplemental Figure 3 and Patient A, image 2, Supplemental Figure 4) and Lesion C (KM-low) (Patient C, image 9, Supplemental Figure 4). The immune cell populations in the matched metastatic immune edge are plotted underneath the mIF image to demonstrate intra-patient, inter-lesional heterogeneity in the same patient. Conversely the adaptive immune cell counts were higher in the KMlow TLR (Patient C). The CD4:CD8 ratio differed between KMhigh (2) and KMlow (0.8). The TLR in patient C (KMlow) displayed a marked increase in regulatory T-cells, neutrophils and macrophages compared with the KMhigh TLRs. In the TLRs from the KMhigh primary CRC, the adaptive immune response was reflected in the corresponding matched CRLM IE, whereas in the CRLM of patient C (KMlow), the CD4 and CD8 counts are almost absent.

**Supplemental Table 1:** GPOL 151 cancer-associated gene panel.

| ABCB1 | CSF1R | FGFR4 | KMT2A | PHF6 | SLC22A1 | ZMYM3 |
| --- | --- | --- | --- | --- | --- | --- |
| ABCC2 | CTNNB1 | FLT1 | KRAS | PIK3CA | SLC22A2 |  |
| ABL1 | CYP19A1 | FLT3 | LAMA2 | PIK3R1 | SLC31A1 |  |
| ABL2 | CYP2A6 | FLT4 | LCK | PSMB1 | SLC34A2 |  |
| AKT1 | CYP2B6 | FSTL5 | LTK | PSMB2 | SLC45A3 |  |
| AKT2 | CYP2C19 | GNA11 | MAP2K1 | PSMB5 | SLCO1B1 |  |
| AKT3 | CYP2C9 | GNAQ | MAP2K2 | PSMD1 | SMAD4 |  |
| ALK | CYP2D6 | GNAS | MAP2K4 | PSMD2 | SMARCA4 |  |
| APC | DDR1 | GSTP1 | MAP3K1 | PTCH1 | SMARCB1 |  |
| ASXL1 | DDR2 | H3F3A | MAPK1 | PTEN | SMO |  |
| ATM | DDX3X | HNF1A | MED13 | PTPN11 | SNCAIP |  |
| ATRX | DNMT3A | HRAS | MET | RAF1 | SOS1 |  |
| BRAF | DPYD | IDH1 | MLH1 | RARA | SPRED1 |  |
| BRCA1 | EGFR | IDH2 | MPL | RARB | SRC |  |
| BRCA2 | ERBB2 | IKZF1 | MST1R | RARG | STK11 |  |
| CBL | ERBB3 | IL2RA | MTOR | RB1 | SUFU |  |
| CDA | ERBB4 | IL2RB | MYC | RET | TAS2R38 |  |
| CDH1 | ERG | IL2RG | MYD88 | ROS1 | TET2 |  |
| CDKN2A | ESR1 | INPP4B | NELL2 | RPS6KB1 | TP53 |  |
| CDKN2B | ESR2 | JAK1 | NF1 | RUNX1 | TRRAP |  |
| CEBPA | EZH2 | JAK2 | NOTCH1 | RXRA | TYK2 |  |
| CHD7 | FBXW7 | JAK3 | NPM1 | RXRB | UGT1A1 |  |
| CHIC2 | FGFR1 | KDM6A | NRAS | RXRG | VHL |  |
| CREBBP | FGFR2 | KDR | PDGFRA | SHH | WT1 |  |
| CRLF2 | FGFR3 | KIT | PDGFRB | SHOC2 | YES1 |  |

**Supplemental Table 2:** The cycling conditions of the PCR plate during the library preparation. Protocol provided by Nanostring.

| Step | Temp. | Time | Cycles |
| --- | --- | --- | --- |
| UDG incubation | 37°C | 30 min | 1 X |
| UDG deactivation | 50°C | 10 min | 1 X |
| Initial denaturation | 95°C | 3 min | 1 X |
| Denaturation | 95°C | 15 sec | 18 X |
| Anneal | 65°C | 60 sec |  |
| Extend | 68°C | 30 sec |  |
| Final extension | 68°C | 5 min | 1 X |
| Hold | 4°C | ∞ | 1 X |

## REFERENCES

1. Dunne DF, Jones RP, Malik HZ, Fenwick SW, Poston GJ. Surgical management of colorectal liver metastases: a European perspective. Hepatic Oncol. 2014 Jan;1(1):121–33.

2. Smith JJ, D’Angelica MI. Surgical management of hepatic metastases of colorectal cancer. Hematol Oncol Clin North Am. 2015 Feb;29(1):61–84.

3. Dhir M, Sasson AR. Surgical Management of Liver Metastases From Colorectal Cancer. J Oncol Pract. 2016 Jan 1;12(1):33–9.

4. Fong Y, Fortner J, Sun RL, Brennan MF, Blumgart LH. Clinical score for predicting recurrence after hepatic resection for metastatic colorectal cancer: analysis of 1001 consecutive cases. Ann Surg. 1999 Sep;230(3):309–18; discussion 318-321.

5. Steele CW, Whittle T, Smith JJ. Review: KRAS mutations are influential in driving hepatic metastases and predicting outcome in colorectal cancer. Chin Clin Oncol. 2019 Oct;8(5):53.

6. Tsilimigras DI, Ntanasis-Stathopoulos I, Bagante F, Moris D, Cloyd J, Spartalis E, et al. Clinical significance and prognostic relevance of KRAS, BRAF, PI3K and TP53 genetic mutation analysis for resectable and unresectable colorectal liver metastases: A systematic review of the current evidence. Surg Oncol. 2018 Jun;27(2):280–8.

7. Datta J, Smith JJ, Chatila WK, McAuliffe JC, Kandoth C, Vakiani E, et al. Coaltered Ras/B-raf and TP53 Is Associated with Extremes of Survivorship and Distinct Patterns of Metastasis in Patients with Metastatic Colorectal Cancer. Clin Cancer Res Off J Am Assoc Cancer Res. 2020 Mar 1;26(5):1077–85.

8. Dekker E, Tanis PJ, Vleugels JLA, Kasi PM, Wallace MB. Colorectal cancer. The Lancet. 2019 Oct 19;394(10207):1467–80.

9. Galbraith NJ, Wood C, Steele CW. Targeting Metastatic Colorectal Cancer with Immune Oncological Therapies. Cancers. 2021 Jul 16;13(14):3566.

10. Pagès F, Mlecnik B, Marliot F, Bindea G, Ou FS, Bifulco C, et al. International validation of the consensus Immunosco re for the classification of colon cancer: a prognostic and accuracy study. The Lancet. 2018 May;391(10135):2128–39.

11. Baldin P, Van den Eynde M, Mlecnik B, Bindea G, Beniuga G, Carrasco J, et al. Prognostic assessment of resected colorectal liver metastases integrating pathological features, *RAS* mutation and Immunoscore. J Pathol Clin Res. 2021 Jan;7(1):27–41.

12. Klintrup K, Mäkinen JM, Kauppila S, Väre PO, Melkko J, Tuominen H, et al. Inflammation and prognosis in colorectal cancer. Eur J Cancer. 2005 Nov 1;41(17):2645–54.

13. Mesker WE, Junggeburt JMC, Szuhai K, de Heer P, Morreau H, Tanke HJ, et al. The carcinoma-stromal ratio of colon carcinoma is an independent factor for survival compared to lymph node status and tumor stage. Cell Oncol Off J Int Soc Cell Oncol. 2007;29(5):387–98.

14. Guinney J, Dienstmann R, Wang X, de Reyniès A, Schlicker A, Soneson C, et al. The consensus molecular subtypes of colorectal cancer. Nat Med. 2015 Nov;21(11):1350–6.

15. Pitroda SP, Khodarev NN, Huang L, Uppal A, Wightman SC, Ganai S, et al. Integrated molecular subtyping defines a curable oligometastatic state in colorectal liver metastasis. Nat Commun. 2018 Dec;9(1):1793.

16. Dunne PD, McArt DG, Bradley CA, O’Reily PG, Barrett HL, Cummins R, et al. Challenging the Cancer Molecular Stratification Dogma: Intratumoral Heterogeneity Undermines Consensus Molecular Subtypes and Potential Diagnostic Value in Colorectal Cancer. Clin Cancer Res. 2016 Aug 15;22(16):4095–104.

17. Fisher NC, Byrne RM, Leslie H, Wood C, Legrini A, Cameron AJ, et al. Biological misinterpretation of transcriptional signatures in tumour samples can unknowingly undermine mechanistic understanding and faithful alignment with preclinical data. Clin Cancer Res Off J Am Assoc Cancer Res. 2022 Jul 6;ccr.22.1102.

18. Pelka K, Hofree M, Chen JH, Sarkizova S, Pirl JD, Jorgji V, et al. Spatially organized multicellular immune hubs in human colorectal cancer. Cell. 2021 Sep;184(18):4734–4752.e20.

19. Wu Y, Yang S, Ma J, Chen Z, Song G, Rao D, et al. Spatiotemporal Immune Landscape of Colorectal Cancer Liver Metastasis at Single-Cell Level. Cancer Discov [Internet]. 2021 Jan 1 [cited 2021 Oct 1]; Available from: https://cancerdiscovery.aacrjournals.org/content/early/2021/08/19/2159-8290.CD-21-0316

20. Merritt CR, Ong GT, Church SE, Barker K, Danaher P, Geiss G, et al. Multiplex digital spatial profiling of proteins and RNA in fixed tissue. Nat Biotechnol. 2020 May;38(5):586–99.

21. Park JH, McMillan DC, Powell AG, Richards CH, Horgan PG, Edwards J, et al. Evaluation of a Tumor Microenvironment-Based Prognostic Score in Primary Operable Colorectal Cancer. Clin Cancer Res. 2015 Feb 15;21(4):882–8.

22. Patel M, Pennel KAF, Quinn JA, Hood H, Chang DK, Biankin AV, et al. Spatial expression of IKK-alpha is associated with a differential mutational landscape and survival in primary colorectal cancer. Br J Cancer [Internet]. 2022 Feb 16 [cited 2022 Apr 15]; Available from: https://www.nature.com/articles/s41416-022-01729-2

23. Mayakonda A, Lin DC, Assenov Y, Plass C, Koeffler HP. Maftools: efficient and comprehensive analysis of somatic variants in cancer. Genome Res. 2018 Nov;28(11):1747–56.

24. Bankhead P, Loughrey MB, Fernández JA, Dombrowski Y, McArt DG, Dunne PD, et al. QuPath: Open source software for digital pathology image analysis. Sci Rep. 2017 Dec;7(1):16878.

25. Therneau T. A Package for Survival Analysis in R_. R package version 3.2-13 [Internet]. 2021. Available from: https://CRAN.R-project.org/package=survival

26. Kassambara A, Kosinski M, Biecek P. survminer: Drawing Survival Curves using “ggplot2”. R package version 0.4.9. [Internet]. Available from: https://CRAN.R-project.org/package=survminer

27. Harrison E, Drake T, Ots R. finalfit: Quickly Create Elegant Regression Results Tables and Plots when Modelling. R package version 1.0.3. [Internet]. 2021. Available from: https://CRAN.R-project.org/package=finalfit

28. Harrell FE. Hmisc: Harrell Miscellaneous. R package version 4.5-0. [Internet]. 2021. Available from: https://CRAN.R-project.org/package=Hmisc

29. Wickham H, Chang W, Henry L, Pedersen TL, Takahashi K, Wilke C, et al. ggplot2: Create Elegant Data Visualisations Using the Grammar of Graphics [Internet]. 2021 [cited 2021 Oct 7]. Available from: https://CRAN.R-project.org/package=ggplot2

30. Robinson MD, McCarthy DJ, Smyth GK. edgeR: a Bioconductor package for differential expression analysis of digital gene expression data. Bioinformatics. 2010 Jan 1;26(1):139–40.

31. Gu Z, Eils R, Schlesner M. Complex heatmaps reveal patterns and correlations in multidimensional genomic data. Bioinforma Oxf Engl. 2016 Sep 15;32(18):2847–9.

32. Korotkevich G, Sukhov V, Budin N, Shpak B, Artyomov MN, Sergushichev A. Fast gene set enrichment analysis [Internet]. Bioinformatics; 2016 Jun [cited 2022 Feb 4]. Available from: http://biorxiv.org/lookup/doi/10.1101/060012

33. Hänzelmann S, Castelo R, Guinney J. GSVA: gene set variation analysis for microarray and RNA-Seq data. BMC Bioinformatics. 2013 Dec;14(1):7.

34. Wu T, Hu E, Xu S, Chen M, Guo P, Dai Z, et al. clusterProfiler 4.0: A universal enrichment tool for interpreting omics data. The Innovation. 2021 Aug 28;2(3):100141.

35. Yu G, He QY. ReactomePA: an R/Bioconductor package for reactome pathway analysis and visualization. Mol Biosyst. 2016 Jan 26;12(2):477–9.

36. Danaher P. SpatialDecon: Deconvolution of mixed cells from spatial and/or bulk gene expression data. 2021.

37. Jackstadt R, van Hooff SR, Leach JD, Cortes-Lavaud X, Lohuis JO, Ridgway RA, et al. Epithelial NOTCH Signaling Rewires the Tumor Microenvironment of Colorectal Cancer to Drive Poor-Prognosis Subtypes and Metastasis. Cancer Cell. 2019 Sep;36(3):319–336.e7.

38. Van den Eynde M, Mlecnik B, Bindea G, Fredriksen T, Church SE, Lafontaine L, et al. The Link between the Multiverse of Immune Microenvironments in Metastases and the Survival of Colorectal Cancer Patients. Cancer Cell. 2018 Dec 10;34(6):1012–1026.e3.

39. Lin HC, Shao Q, Liang JY, Wang Y, Zhang HZ, Yuan YF, et al. Primary tumor immune score fails to predict the prognosis of colorectal cancer liver metastases after hepatectomy in Chinese populations. Ann Transl Med. 2021 Feb;9(4):310–310.

40. Huang M, Wu R, Chen L, Peng Q, Li S, Zhang Y, et al. S100A9 Regulates MDSCs-Mediated Immune Suppression via the RAGE and TLR4 Signaling Pathways in Colorectal Carcinoma. Front Immunol. 2019 Sep 18;10:2243.

41. Wang C, Gu Y, Zhang K, Xie K, Zhu M, Dai N, et al. Systematic identification of genes with a cancer-testis expression pattern in 19 cancer types. Nat Commun. 2016 Jan 27;7(1):10499.

42. Schürch CM, Bhate SS, Barlow GL, Phillips DJ, Noti L, Zlobec I, et al. Coordinated Cellular Neighborhoods Orchestrate Antitumoral Immunity at the Colorectal Cancer Invasive Front. Cell. 2020 Sep 3;182(5):1341–1359.e19.

43. Park JH, Richards CH, McMillan DC, Horgan PG, Roxburgh CSD. The relationship between tumour stroma percentage, the tumour microenvironment and survival in patients with primary operable colorectal cancer. Ann Oncol Off J Eur Soc Med Oncol. 2014 Mar;25(3):644–51.

44. Huijbers A, Tollenaar R a. EM, v Pelt GW, Zeestraten ECM, Dutton S, McConkey CC, et al. The proportion of tumor-stroma as a strong prognosticator for stage II and III colon cancer patients: validation in the VICTOR trial. Ann Oncol Off J Eur Soc Med Oncol. 2013 Jan;24(1):179–85.

45. Ståhl PL, Salmén F, Vickovic S, Lundmark A, Navarro JF, Magnusson J, et al. Visualization and analysis of gene expression in tissue sections by spatial transcriptomics. Science. 2016 Jul 1;353(6294):78–82.

46. Hoang ML, Kriner M, Zhou Z, Norgaard Z, Sorg K, Merritt C, et al. Abstract 1364: Spatially-resolved *in situ* expression profiling using the GeoMx^TM^ Cancer Transcriptome Atlas panel in FFPE tissue. In: Molecular and Cellular Biology / Genetics [Internet]. American Association for Cancer Research; 2020 [cited 2021 Nov 25]. p. 1364–1364. Available from: http://cancerres.aacrjournals.org/lookup/doi/10.1158/1538-7445.AM2020-1364

